# Multimodal cell tracking from systemic administration to tumour growth by combining gold nanorods and reporter genes

**DOI:** 10.1101/199836

**Authors:** Joan Comenge, Jack Sharkey, Oihane Fragueiro, Bettina Wilm, Mathias Brust, Patricia Murray, Raphaël Lévy, Antonius Plagge

## Abstract

Understanding the fate of exogenous cells after implantation is important for clinical applications. Preclinical studies allow imaging of cell location and survival. Labelling with nanoparticles enables high sensitivity detection, but cell division and cell death cause signal dilution and false positives. By contrast, genetic reporter signals are amplified by cell division. Here we characterise lentivirus-based bi-cistronic reporter gene vectors and silica-coated gold nanorods (GNRs) as synergistic tools for cell labelling and tracking. Co-expression of the bioluminescence reporter luciferase and the optoacoustic reporter near-infrared fluorescent protein iRFP720 enabled cell tracking over time in mice. Multispectral optoacoustic tomography (MSOT) showed immediate biodistribution of GNR-labelled cells after intracardiac injection and successive clearance of GNRs (day 1-15) with high resolution, while optoacoustic iRFP720 detection indicated tumour growth (day 10-40). This multimodal cell tracking approach could be applied widely for cancer and regenerative medicine research to monitor short- and long-term biodistribution, tumour formation and metastasis.

## Introduction

Non-invasive optical imaging methods for preclinical *in vivo* research include bioluminescence (BLI) and fluorescence as well as photoacoustic/optoacoustic tomography, a technology that has only been developed recently ^1–3^. These imaging modalities have enabled great progress in the tracking of labelled cells longitudinally over time in animal models of disease, which has become especially relevant for cancer research and cell-based regenerative medicine therapies ^4–6^. The resolution and sensitivity of optical imaging in animals is limited by autofluorescence, absorption and scattering of excitation and/or emission light, especially in deep tissues. The optimal window for *in vivo* optical imaging lies in the near infrared (NIR) spectrum (~650-900 nm), since absorption through the main endogenous chromophores (oxy-haemoglobin, deoxy-haemoglobin, melanin, water and lipids) are minimal in this spectral range ^3^. For permanent cell labelling and tracking, genetic modification with reporter genes is the method of choice, although fluorescent tags and nanoparticles have been developed recently for sensitive short-term cell tracking over a period of a few cell divisions ^7,8^.

Using luciferase reporter genes, bioluminescence constitutes the most sensitive optical modality due to its excellent signal-to-noise ratio, as light emission only occurs in the presence of a functional enzyme and its required co-factors. Firefly luciferase has become the most widely used reporter as its substrates, D-luciferin or CycLuc1 ^9^, are very well tolerated by animals and, compared to other luciferases, its peak light emission at around 562 nm is closest to the infrared window for *in vivo* imaging ^4^. Although highly sensitive *in vivo* cell tracking via bioluminescence imaging of firefly luciferase is well established ^4,10^, this modality provides poor information about the spatial localisation of cells. Fluorescence has recently gained importance for animal imaging, since novel near-infrared fluorescent proteins (iRFPs) were developed from bacterial phytochrome photoreceptors ^11,12^. Similar to bioluminescence imaging, fluorescence only allows limited spatial resolution due to the high scattering coefficient of photons in tissues.

On the other hand, photoacoustic imaging is based on the generation of ultrasound waves after absorption of light emitted by a pulsed laser. The sound waves are well transmitted in fluid media and less prone to scattering through tissues than emitted light. In fact, acoustic scattering is three orders of magnitude less than photon scattering ^13^, which overcomes deep tissue spatial resolution drawbacks of other optical-based imaging technologies. Interestingly, some iRFPs, such as iRFP720, have an absorption profile in the NIR window, thus enabling their use as reporter genes for photoacoustic imaging, and allowing deep tissue imaging and tumour monitoring in mice ^1,14^. For example, new iRFPs have been proven to be useful genetic photoacoustic reporters in mammary gland and brain tumour monitoring, which establishes them as dual-modality imaging probes ^1,15,16^. In addition, in multispectral optoacoustic tomography (MSOT), a rapid multiwavelength excitation allows the distinction between different absorbers simultaneously after applying multispectral unmixing algorithms ^17^. Hence, a number of endogenous (e.g. deoxy- and oxyhaemoglobin) or exogenously introduced targets can be imaged *in vivo* via MSOT, including fluorescent proteins and nanoparticles.

Since light absorption triggers the photoacoustic response, contrast agents with a high molar extinction coefficient facilitate their distinction from endogenous chromophores. In this regard, GNRs are an attractive choice since the extinction coefficients are in the range of 4 to 5.5 × 10^9^ M^−1^cm^−1^ for longitudinal surface plasmon between 728 and 845 nm^18^, i.e. orders of magnitude higher than endogenous absorbers or other types of contrast agents (e.g. AlexaFluor750 and iRFP720 coefficients are 2.5 × 10^5^ and 9.6 × 10^4^ M^−1^cm^−1^, respectively ^12,19^). We recently demonstrated how, after coating GNRs with a 35 nm silica shell, the optical properties of GNRs are maintained even in harsh environments such as the endosomal vesicles^7^. This enabled very sensitive photoacoustic imaging of labelled cells, pushing the limits of detection into a range useful for biologically-relevant applications including cell tracking. However, cell division and cell death result in a decrease of GNR concentration and therefore a decrease of photoacoustic intensity. Thus, for long-term photoacoustic tracking of cells, adequate reporter genes are required.

Here we characterise newly generated lentivirus-based bi-cistronic reporter gene vectors and silica-coated GNRs as synergistic tools for co-labelling of cells and tracking in multiple imaging modalities at different stages after cell administration. The GNR-860 offered an excellent photoacoustic sensitivity that enabled detection of a very low number of cells.

Reporter gene vectors mediated co-expression of the most sensitive bioluminescent reporter, firefly luciferase, and the most red-shifted near-infrared fluorescent protein, iRFP720, which also acted as a photoacoustic reporter in MSOT. Whilst GNR-860 enabled early detection of cells by photoacoustic imaging after systemic injection, reporter genes were needed to monitor any potential tumour growth with the same technique. In addition, co-expression of luciferase allowed bioluminescence imaging. Although this modality does not offer the spatial resolution of photoacoustic imaging, it was used here to continuously monitor cell viability as well as for validating the results of photoacoustic tomography. The combination of the two cell labelling strategies (nanoparticles and genetic labelling) allowed short-term and long-term multimodal imaging of deep tissues with high spatial resolution without compromising sensitivity. It enabled the non-invasive monitoring of the initial cell biodistribution on a sub-organ level as well as the detection of tumour formation from very early stages, thus opening opportunities for gaining a better understanding of stem cell therapies safety required for moving towards clinical translation^20^.

## Methods

### Reagents

The following chemicals were purchased from Sigma-Aldrich (Gillingham, UK). HAuCl_4_ ·3H_2_O (>99.0%), NaBH_4_ (>99.99%), AgNO_3_(99.0%), hexadecyltrimethylammonium bromide (CTAB, >99%), L-ascorbic acid (reagent grade), tetraethyl orthosilicate (TEOS, 99.999%), O-[2-(3-Mercaptopropionylamino)ethyl]-O′-methylpolyethylene glycol (mPEG-SH, MW: 5000 Da), 5-bromosalicylic acid. Monocarboxy (1-mercaptoundec-11-yl) hexaethylene glycol (PEG-COOH, MW 526.73 Da) was obtained from Prochimia (Sopot, Poland). Dulbecco's modified eagle's medium (DMEM), phosphate buffered saline (PBS), penicillin-streptomycin, biliverdin and polybrene were also obtained from Sigma Aldrich. Foetal bovine serum (FBS) was purchased from Life-Technologies. D-Luciferin was purchased from Promega (Southampton, UK) and CycLuc1 from Glixx Laboratories (Southborough, USA).

### Gold nanorod synthesis

GNR synthesis were based on previously published protocols ^21,22^. First, seeds were prepared by adding 0.6 mL of ice-cold NaBH_4_ (0.01 M) to a mixture of 5 mL CTAB (0.2 M) and 5 mL of HAuCl_4_ (0.5 mM) under vigorous stirring. The growth solution for GNR-710 was prepared by mixing 50 mL of CTAB (0.2M), 1.8 mL of AgNO_3_ (4 mM), 50 mL of HAuCl_4_ (1 mM) and 0.8 mL of ascorbic acid (0.1 M). Finally, 0.2 mL of freshly synthesised seeds was added to the growth solution. The reaction was kept in a water bath at 28°C for 3 h. The growth solution for GNR-860 was prepared by adding 440 mg of 5-bromosalicylic acid to 100 mL of CTAB (0.1 M) and heat up the mixture to 60°C. After cooling down the solution to 30°C, 4.8 mL of AgNO_3_ (4 mM) were added and left undisturbed for 15 min. Then, 50 mL of HAuCl_4_ (1 mM) were added and left under low stirring for 15 min at 30°C. Afterwards, 256 |aL of ascorbic acid was added under vigorous agitation for 30 s. Finally, 160 μL of seeds were added and the solution was kept at 28°C for 12 h.

GNRs were visualized using a Tecnai G3 Spirit transmission electron microscope (TEM) at 120 keV. Formvar/carbon-coated 200 mesh copper grid (TAAB) were dipped in a solution of the GNRs of interest and left to dry in air. More than 100 GNRs were considered for image analysis.

### Lentivirus vectors

Lentivirus vectors were constructed on the pHIV-Luciferase backbone (a gift from Bryan Welm, Addgene plasmid # 21375). The iRFP720 ORF was obtained from plasmid piRFP720-N1 (a gift from Vladislav Verkhusha, Addgene plasmid # 45461)^12^ and cloned as an EcoRI/XbaI fragment upstream of the IRES element of pHIV-Luciferase, resulting in the pHIV-iRFP720-IRES-Luc vector. To achieve a more efficient translation of the second ORF, the IRES element was replaced with an E2A peptide motif ^23,24^ using a two-step PCR protocol to create the pHIV-iRFP720-E2A-Luc plasmid. The complete sequence of the E2A plasmid version has been submitted to the NCBI GenBank database (accession number: MF693179). Lentivirus particles were produced in HEK293 cells via calcium phosphate cotransfection of the reporter gene plasmid, packaging plasmid psPAX2 and envelope plasmid pMD2.G (both a gift of Didier Trono, Addgene plasmids #12260 and #12259, respectively) as described ^25^. Virus-containing cell culture supernatant was collected three days post-transfection, centrifuged at 500 g and filtered through 0.45 μm pores.

### Mouse mesenchymal stem/stromal cell (mMSC) line culture, labelling and immunofluorescence

We chose the mouse MSC line D1 ORL UVA [D1](ATCC^®^ CRL-12424^TM^) for labelling and tracking, since these cells have the potential to differentiate into mesenchymal derivatives, but can occasionally also form tumours as previously shown^7,26^. They were grown in DMEM supplemented with 10% FBS and 2 mM L-Glutamine. Cells were transfected with purified reporter gene virus particles for 24 h. iRFP720-expressing cells were sorted using a BD FACSAria III with red laser and APC-Cy^TM^7 filter. Cells expressing high levels of iRFP720 were selected and maintained for all further experiments. Before IVIS imaging of cells, biliverdin was added to the culture medium for 24 h to increase its concentration beyond levels present in FBS and to improve its uptake by cells for incorporation into iRFP720 as a chromophore. For analysis of reporter gene expression by immunofluorescence, mMSCs were grown on glass cover slips, fixed with 4% paraformaldehyde/PBS, washed with PBS, blocked and stained with an anti-firefly luciferase antibody (Abcam, Cambridge, UK, ab21176, diluted in 1:500 in PBS/10% donkey serum/0.25% Triton-X100) and a donkey-anti-rabbit-AlexaFluor488 secondary antibody (Molecular Probes, A21206). Fluorescence of iRFP720 was assessed directly using a Zeiss LSM510 Multiphoton microscope.

GNR-labelling was performed as previously published ^7^. Briefly, cells were treated with cell medium containing GNRs at the concentration / optical density indicated in the main text (always 79% cell medium, 20% GNRs in water, 1% penicillin-streptomycin). Then, cells were dissociated with trypsin, resuspended in fresh medium, washed twice with PBS, and counted using an automated cell counter (TC10, BioRad, Watford, UK).

### Cell viability

Cell viability was assessed with Cell Titer Glo ATP Assay (Promega). Cells were labelled as described above by seeding 10^4^ mMSCs cells in 96-well plates. Cells were incubated with GNRs-710 and GNR-860 at a final O.D. in media=1, 2, 3.2, and 4 (80% media and 20% GNRs solution in each case) for 24 h. After labelling, cells were washed 3 times with PBS. 50 μL of medium were added to each well and then 25 μL of the ATP reagent was added. The plate was mixed in an orbital shaker and, after 10 min, the contents of the plate were transferred to white, opaque, 96-well plates and the luminescence measured with a plate reader (Fluostar Omega, BMG Labtech, Aylesbury, UK). Each condition was assessed in triplicate and results are given as % ± SD in respect to cells that were incubated with no GNRs as described above.

### Preparation of cells for TEM

Cells were fixed with a solution containing 1% paraformaldehyde and 3% glutaraldehyde in 0.1 M cacodylate buffer (pH 7.4). Then, cells were incubated with a reduced osmium staining solution, containing 2% OsO4 and 1.5% K_4_[Fe(CN)_6_], for 1 hour. This was followed by a second 1 h osmium staining (2% OsO_4_) step and overnight staining with 1% uranyl acetate. Cells were washed with water for 3 min, 3 times after every staining step. Samples were then dehydrated in graded ethanol (30%, 50%, 70%, 90% and 2× 100%) for 5 min each. Finally, samples were infiltrated with medium TAAB resin 812 and embedded within the same resin. The resin was cured for 48 h at 60°C. Finally, ultrathin sections of 350 μm × 350 μm × 74 nm were cut and placed in 200-mesh Formvar/Carbon filmed grids. They were post-stained with uranyl acetate (4% UA in a 50:50 ethanol/water solution) and Reynolds lead citrate before TEM imaging.

### Animals

8-10 week old female SCID hairless outbred (SHO) mice (Charles River, Margate, UK) were housed in individually ventilated cages at a 12 h light/dark cycle, with *ad libitum* access to food and water. Experimental animal protocols were performed in accordance with the guidelines under the Animals (Scientific Procedures) Act 1986 (licence PPL70/8741) and approved by the University of Liverpool Animal Welfare and Ethical Review Body. The tumour burden was monitored and kept within recommended limits in accordance with guidelines for the welfare and use of animals in cancer research^27^. Experiments are reported in line with the ARRIVE guidelines. These experiments aimed at evaluating the potential of MSOT to track cells over the short and long-term. The number of animals was chosen so that a range of tumour positions and sizes could be observed (Table 1). Image quantification is presented to demonstrate the information that can be extracted from this approach. In each case, the numbers correspond to the particular animal presented. The data for these exemplar animals as well as for the other animals are available in the data repository Zenodo ^28^.

**Table 1.**
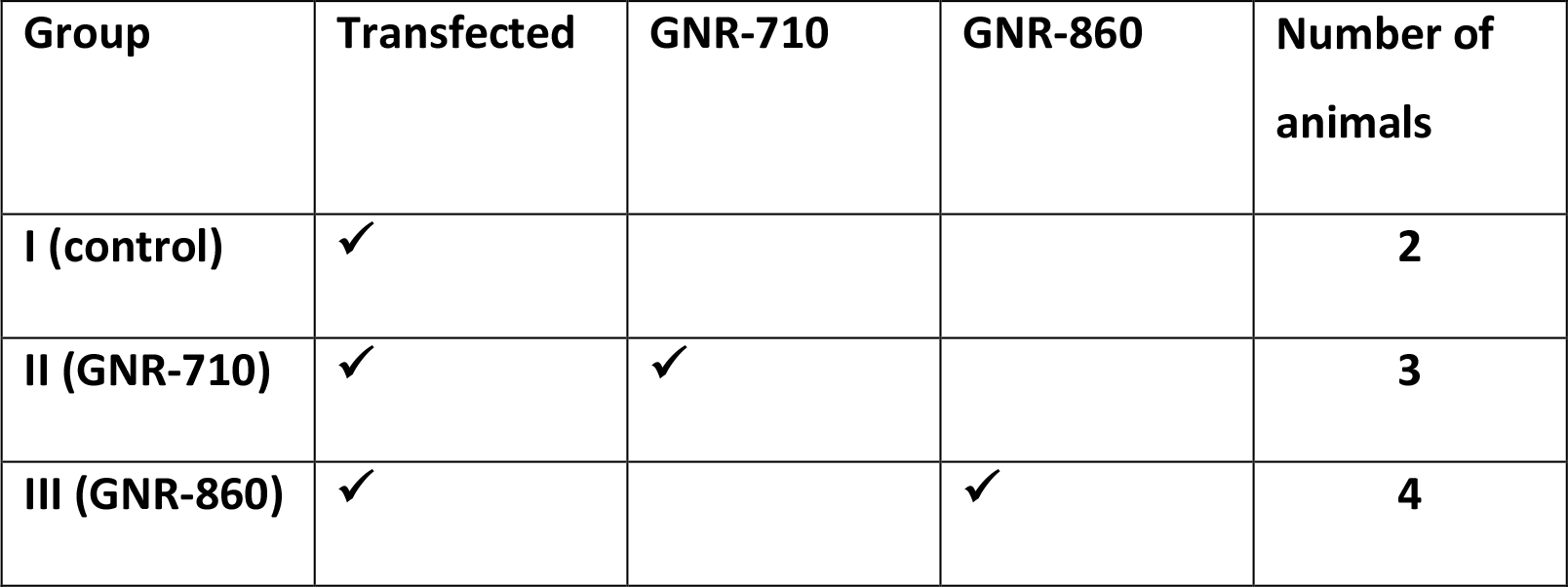
Summary of mice used and analysed for the experiment.

### Intracardiac injection and imaging

Transfected cells were labelled with GNR-710 and GNR-860 at O.D.=2.4 as detailed above. 1.5 × 10^6^ cells in 100 μL PBS were prepared for intracardiac injection as described below. All cell injection and imaging procedures of mice were carried out under general isoflurane/oxygen anaesthesia.

The imaging routine at day 0 was composed of the following sequence of steps: 1) a baseline MSOT scan; 2) the ultrasound-guided (Prospect imaging system, S-Sharp, New Taipei City, Taiwan) intracardiac injection of cells into the left ventricle; 3) a second MSOT scan after 10 min for acclimatisation; 4) BLI, for which the mouse received an intraperitoneal injection of D-luciferin (approximately 150 μg/g body weight) 15 min before whole body imaging using an IVIS Spectrum system (Perkin Elmer, Seer Green, UK). All data were analysed with Living Image (Perkin Elmer) and data are displayed in radiance units. On subsequent days the MSOT scan and BLI were repeated in the same manner.

For intracardiac injection of cells, mice were positioned supine on a heated platform. Fur around the chest area was removed using depilatory cream and limbs were taped down to keep the mouse position fixed. Ultrasound gel was applied liberally to the chest area and the ultrasound transducer (Prospect imaging system, S-Sharp) was positioned above the chest so the long axis view of the left ventricle was visible. 100 μl of cell suspension was drawn up into an insulin syringe (29 G) and, using the ultrasound image as guidance, was inserted into the left ventricle of the heart. Cell suspension was then administered slowly over a period of approximately 30 s.

For optoacoustic imaging, an MSOT inVision 256-TF small animal imaging system (iThera Medical GmbH, Munich, Germany) was used ^29^. After acclimatisation for 10 min inside the water bath, a whole-body scan was performed on the mouse with 1 mm steps and the following wavelengths for acquisition: 660, 670, 680, 690, 700, 705, 710, 715, 725, 735, 750, 765, 780, 795, 810, 820, 830, 840, 850, 860, 870, 880, 890, 900, 910, 920, 930, 940, 1025, 1050, 1075, and 1100 nm. Heavy water was used in the water bath due to its low absorbance at wavelengths >910 nm (contrary to regular water). Linear-mode-based reconstruction and guided ICA multispectral processing were applied using viewMSOT software v3.6 (iThera Medical GmbH).

## Results and discussion

### Gold nanorod synthesis and cell labelling

Two types of GNRs with distinct optical spectra were prepared following previously published protocols ^21,22^. The first batch consisted of GNRs with a longitudinal surface plasmon resonance (LSPR) peak at 710 nm (GNR-710). Their core size was 60.0 ± 7.5 nm by 25.7 ± 3.7 nm with an average aspect ratio of 2.3 ± 0.3. The second batch had a LSPR peak at 860 nm (GNR-860). In this case, their size was 93.8 ± 9.3 nm by 25.2 ± 4.4 nm with an average aspect ratio of 3.8 ± 0.5) (Figure 1a). Both GNRs were prepared to have LSPR bands in the NIR to favour their detection deep inside tissues.

We have recently demonstrated that a 35 nm silica shell is needed to minimise plasmon coupling between GNRs after cell uptake ^7^. This enables the preservation of intrinsic optical properties resulting in an increased sensitivity for photoacoustic detection. Hence, both GNR batches were coated with a silica shell as previously described. Specifically, GNR-710 and GNRs-860 nm have a silica shell thickness of 38.7 ± 1.9 nm and 35.7 ± 1.6 nm respectively (Figure 1a).

A murine MSC line was selected for labelling and tracking, since these cells have the capability to differentiate into mesenchymal derivatives, but can also form tumours occasionally^7,26^. The cells, which had first been transfected with a reporter gene vector (see below), were incubated with different concentrations of GNR-710 and GNR-860 for 24 hours. To provide a better comparison between GNRs of different sizes, we standardize and report the GNR concentrations by the optical density (O.D.) of the peak, because absorption is what triggers the photoacoustic response. For reference, O.D.=4 corresponds to ~100 pM GNR-710 and ~72.5 pM GNR-860 (calculated by relating the absorbance at 400 nm of GNRs before silica coating to concentrations of molecular gold) ^30^.

To characterize their uptake, cells were incubated with GNRs at 0.D.=3 for 24 h. TEM images of cells show GNRs localised inside vesicles, with silica shells separating the gold cores (Figure 1b, and Supplementary file 1). In line with our previous report, the optical properties of GNRs were largely preserved after cell uptake (Figure 1d). Preservation of the absorbance intensity and the shape of the plasmon band are prerequisite for an optimal photoacoustic detection of GNR-labelled cells since it relies on a spectral deconvolution to differentiate intrinsic absorbers from probes.

To confirm that the cell labelling does not cause any toxicity, cell viability after exposure to a range of GNR concentrations up to O.D.=4 was assessed (Figure 1e). Only at the highest GNR concentration was a slight decrease in cell viability observed: 92.0 ± 2.6% and 91.7 ± 1.6% for GNR-710 and GNR-860, respectively. These results are in agreement with our previous work, in which we showed that silica coated GNRs did not affect cell viability, proliferation or differentiation potential of the mMSCs ^7^.

**Figure 1.**
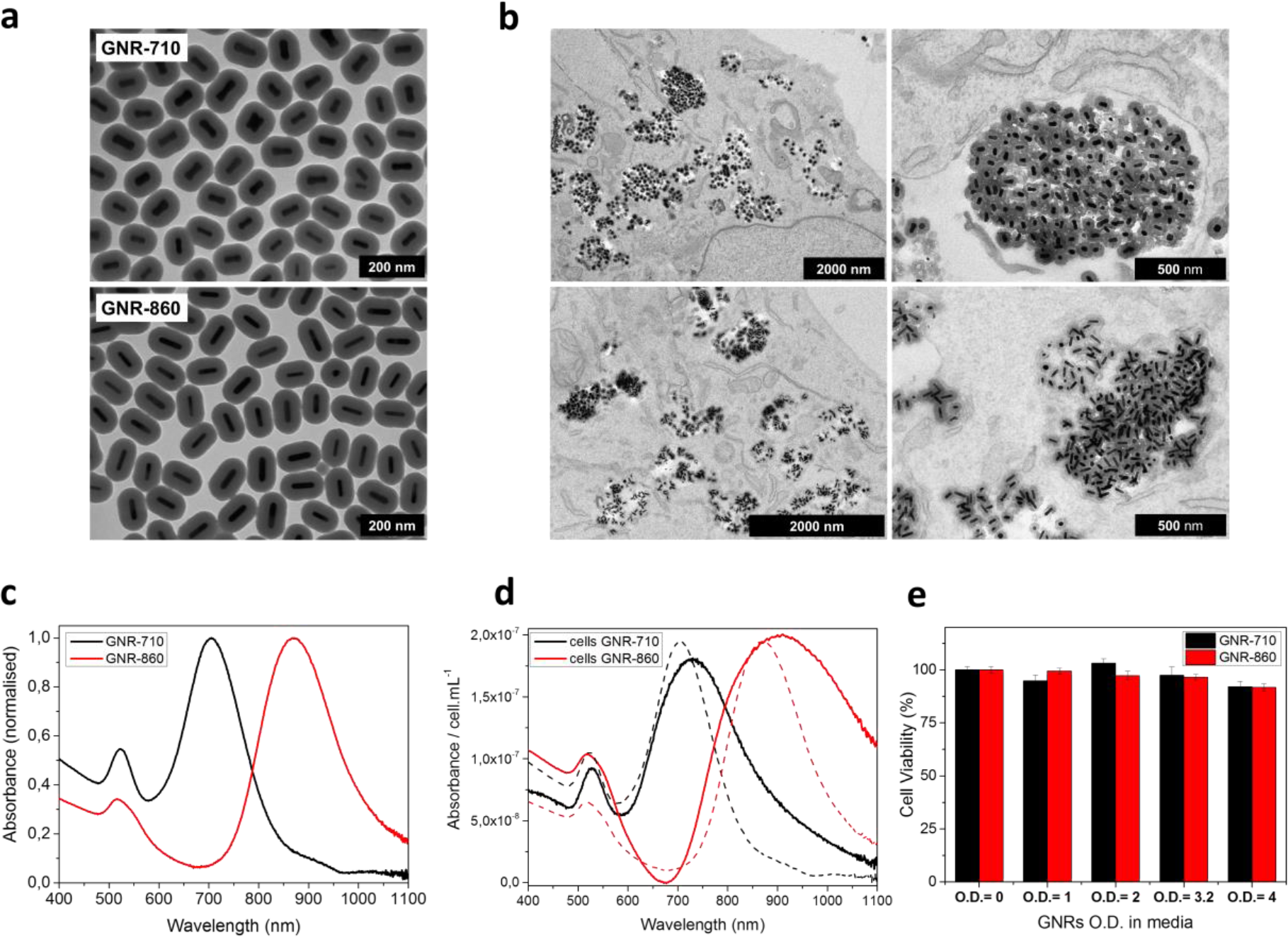
GNR characterization and cell labelling. (a) Representative TEM pictures of GNR-710 (top) and GNR-860 (bottom). (b) TEM pictures of GNR-710 (top panels) and GNR-860 (bottom panels) inside cells. In both cases, the silica shell provides steric hindrance to minimise plasmon coupling in cellular vesicles. (c) Vis-NIR spectrum of GNRs in solution. (d) Vis-NIR spectrum of cells labelled with the corresponding GNRs. Dashed lines correspond to GNRs in solution. (e) Cell viability at different optical densities.

### Generation of iRFP720-Luciferase bi-cistronic lentivirus reporter gene vectors and characterisation of labelled cells

To combine the sensitivity of bioluminescence imaging with the spatial resolution of MSOT, we generated bi-cistronic lentivirus vectors for expression of firefly luciferase and iRFP720 from the general eukaryotic EF1α promoter (Figure 2a). We compared vectors containing either an internal ribosome entry site (IRES) ^31^ or a self-cleaving 2A element ^23 24^, to determine the most efficient method of translating the second open reading frame (ORF; encoding luciferase) from a single mRNA. Following transfection with lentivirus vectors, the mMSCs were selected by FACS for iRFP720 fluorescence, to obtain an IRES-vector and an E2A-vector transfected cell population, respectively, with similar levels of expression of the first ORF protein (iRFP720) (Figure 2b). Co-expression of iRFP720 and luciferase in individual cells was confirmed by immunocytochemistry (Figure 2c), which also indicated that, in contrast to luciferase, iRFP720 was preferentially localised in the nucleus. To compare the levels of expression of luciferase from the IRES-vector and E2A-vector transfected cells, fluorescent and bioluminescent signals were quantified in cell suspensions using an IVIS Spectrum imager. For both types of reporter cells (IRES-vector and E2A-vector transfected), the fluorescence signal intensity for iRFP720 was the same, but the E2A-vector transfected cells showed a 5 times higher luciferase bioluminescence (Figure 2d, e). These data indicate that in the context of our bi-cistronic lentivirus vectors, a superior translation efficiency of the second ORF from a single mRNA was achieved via an E2A element as compared to an IRES element. We therefore used the pHIV-iRFP720-E2A-Luc vector in all further experiments described herein. Proof of principle of the application of this vector for multi-modal imaging was tested in preliminary experiments, in which cells were injected subcutaneously and monitored with bioluminescence, fluorescence and photoacoustic imaging (Supplementary file 2).

**Figure 2.**
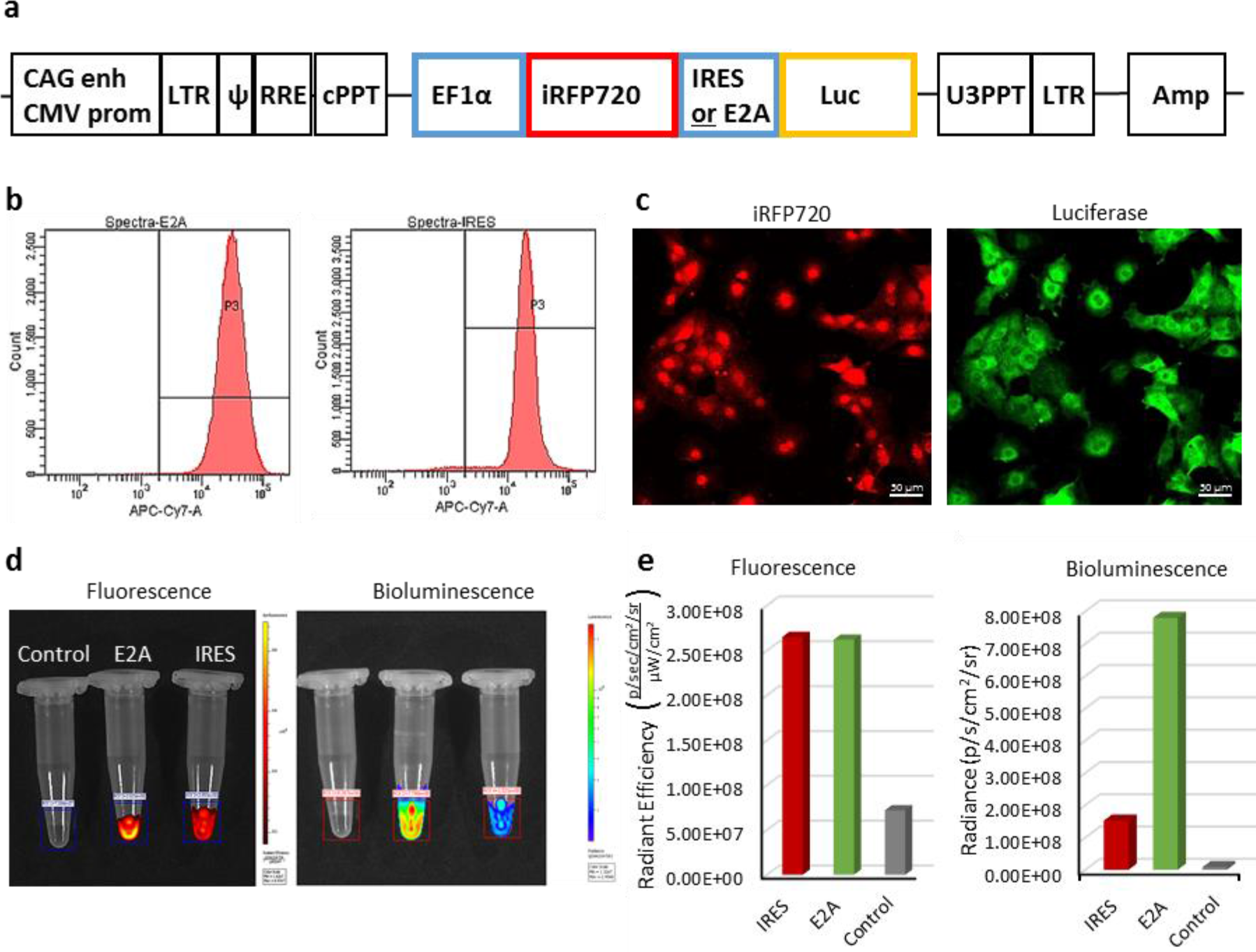
Characterisation of iRFP720-Luciferase bi-cistronic lentivirus reporter gene vectors and transfected cells *in vitro*. a) Scheme of the vectors that mediate co-expression of iRFP720 and firefly luciferase either via IRES- or E2A-mediated mechanisms. b) FACS analyses of iRFP720 fluorescent signal intensity in mMSCs transfected with either the IRES or E2A vector, respectively, indicated similar levels of expression of the first ORF. c) Confocal fluorescence microscopy of transfected cells for co-expression of iRFP720 and luciferase (immunofluorescence). In contrast to Luc, iRFP720 accumulated in the nucleus. d), e) Comparison of fluorescent and bioluminescent signal intensities from IRES and E2A vector-transfected cell populations using IVIS Spectrum imaging. Although both cell suspensions provided very similar levels of fluorescence from the first reporter ORF (iRFP720) in the region of interest (ROI), bioluminescence levels were 5-fold higher in the E2A vector-transfected cell population, indicating a more efficient translation of the second ORF (luciferase) from the E2A element than from the IRES. Fluorescence was measured first, followed by addition of CycLuc1 to the cell suspension and bioluminescence imaging.

### Imaging biodistribution of cells after intracardiac injection

In our previous work ^7^ and in many cell tracking studies, proof of principle of the utility of nanoparticles as contrast agents for photoacoustic cell detection has been obtained by imaging labelled cells implanted locally ^32, 33^. Here, we address the much more challenging case of a systemic injection where cells spread through the body and the signal, therefore, becomes diluted. Specifically, 1.5 × 10^6^ mMSCs transduced with the iRFP720-E2A-Luc lentivirus and labelled with GNRs were administered via ultrasound-guided intracardiac injection into the left ventricle. Left ventricle administration was chosen for cells to enter the arterial circulation rather than the venous. This allows cells to pass through all organs before returning to the right ventricle and subsequently lungs, where they become sequestered in the pulmonary microvasculature ^34,35^. To provide a detailed assessment of cell localisation, we combined photoacoustic tomography imaging at 1 mm steps with bioluminescence detection. While bioluminescence imaging provides excellent sensitivity, its anatomical resolution is poor mainly due to the high scattering of photons by tissue ^36,37^. By contrast, photoacoustic imaging provides superior spatial resolution (150 μm in our system) ^19^. To demonstrate the potential of this multimodal approach to precisely determine the localisation of distributed nanorod-labelled cells, we show the correlation between bioluminescence imaging and MSOT imaging (Figure 3). Cell localisation in the head, liver and kidney regions was clearly observed with both imaging modalities (Figure 3a, b), and the cell presence within organs was confirmed in the photoacoustic scans through the respective regions (Figure 3c). This distribution is in agreement with other, different types of assessments in previous reports ^38^. The presence of cells in the brain region is probably due to cell trapping in small capillaries, as reported also in other studies after intracardiac injection of cells ^39^. It should be noted here that there was no difference in the bioluminescence signal biodistribution when cells with or without gold nanorods were injected, indicating that GNR uptake does not affect the overall behaviour of the cells (Supplementary file 4).

**Figure 3.**
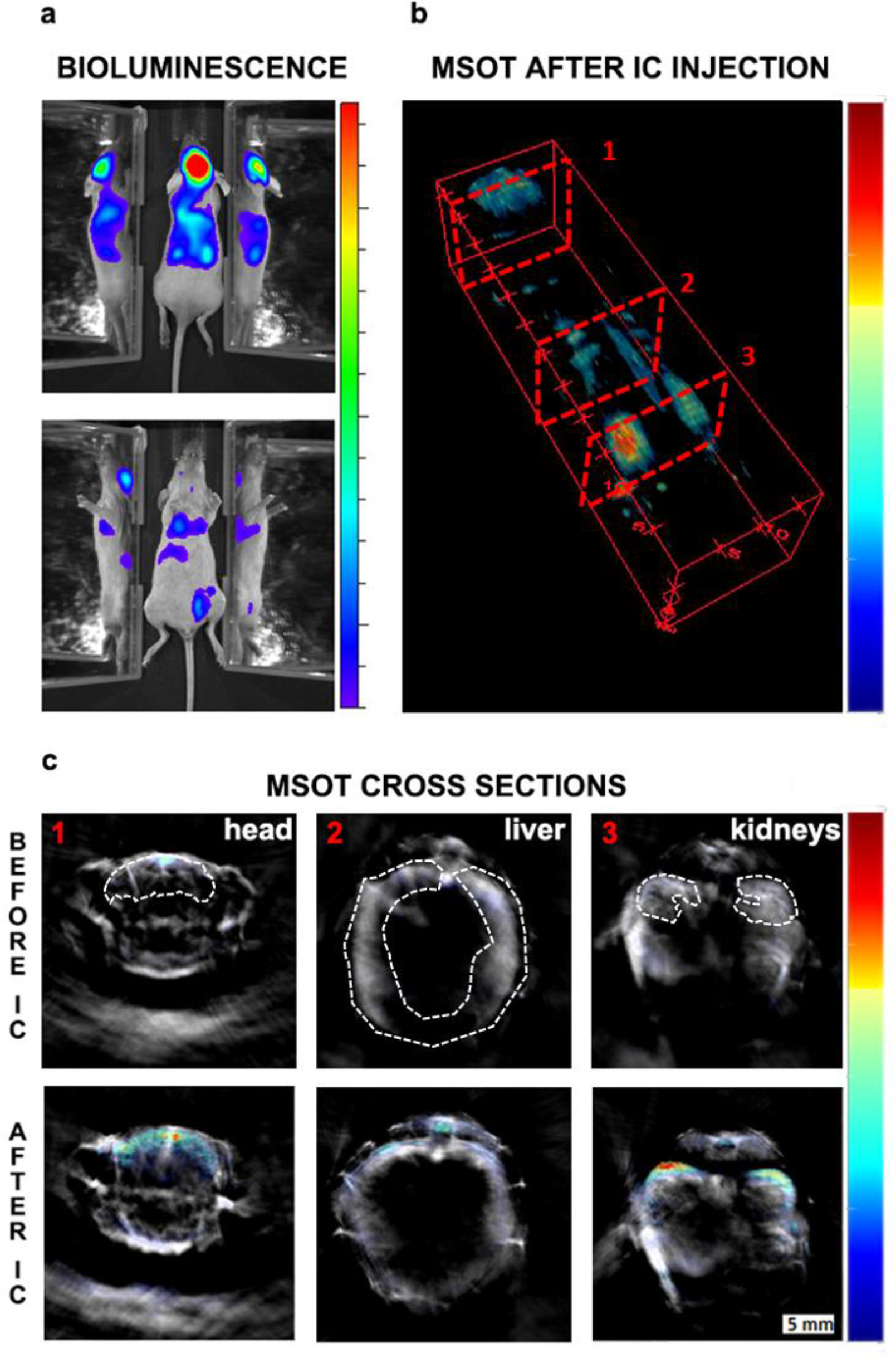
Initial biodistribution of GNR-860 labelled cells after intracardiac administration. (a)Bioluminescence imaging in dorsal (top panel) and ventral (bottom panel) position after cell administration. Mirrors were placed on the side of the mouse to provide a lateral view.(b) MSOT imaging of the same animal: 3D reconstruction for the whole animal showed the correlation between GNR-860 signal with bioluminescence imaging (360° animation available in Supplementary File 3). (c) Cross sections corresponding to the planes shown in (b) before and after the injection of cells. Sub-organ localisation of cells can be observed here (see main text for details and supporting information for the whole set of cross section images). The brain, liver and kidney are outlined. Colour scales are 3.0 × 10^5^ to 1.8 × 10^6^ radiance units (ρ· s^−1^ · cm^−2^ · sr^−1^) in (a) and 0.9 to 21 MSOT intensity (a.u.) in (b) and (c).

To further test the robustness and specificity of the signal, additional controls were performed. First, following the same imaging protocols, we observed a similar biodistribution pattern for mice injected with either GNR-710 or GNR-860 labelled cells (Supplementary files 4, 5). Second, when we applied the unmixing algorithm with the ‘wrong’ spectrum (i.e. the GNR-710 spectrum to an animal injected with GNR-860 labelled cells, and vice versa), no GNR-specific signals were detected in photoacoustic images pre-and post-injection (Figure 4). Third, when the photoacoustic spectra of regions of interest before and after injection of cells were extracted, an increase in the photoacoustic intensity was only observed in the range of wavelengths in which corresponding GNRs have the higher absorbance, while intensity in the other wavelengths remained similar (Supplementary file 6). It has to be noted that in some cases there were regions with an endogenous absorbance similar to GNRs (e.g. food in the intestines has similar spectra as GNR-710), which can lead to misinterpretation (Supplementary file 5). For this reason, a scan was performed before cells were injected, which allowed detection of any potential endogenous interference with the GNR signal. Using a multimodal imaging approach (in this case luminescence and photoacoustic) also assisted in discounting any false positive signals. As expected, the photoacoustic signal provided by iRFP720 expression was not strong enough to detect the cells immediately after injection by means of photoacoustic imaging (Figure 4).

**Figure 4.**
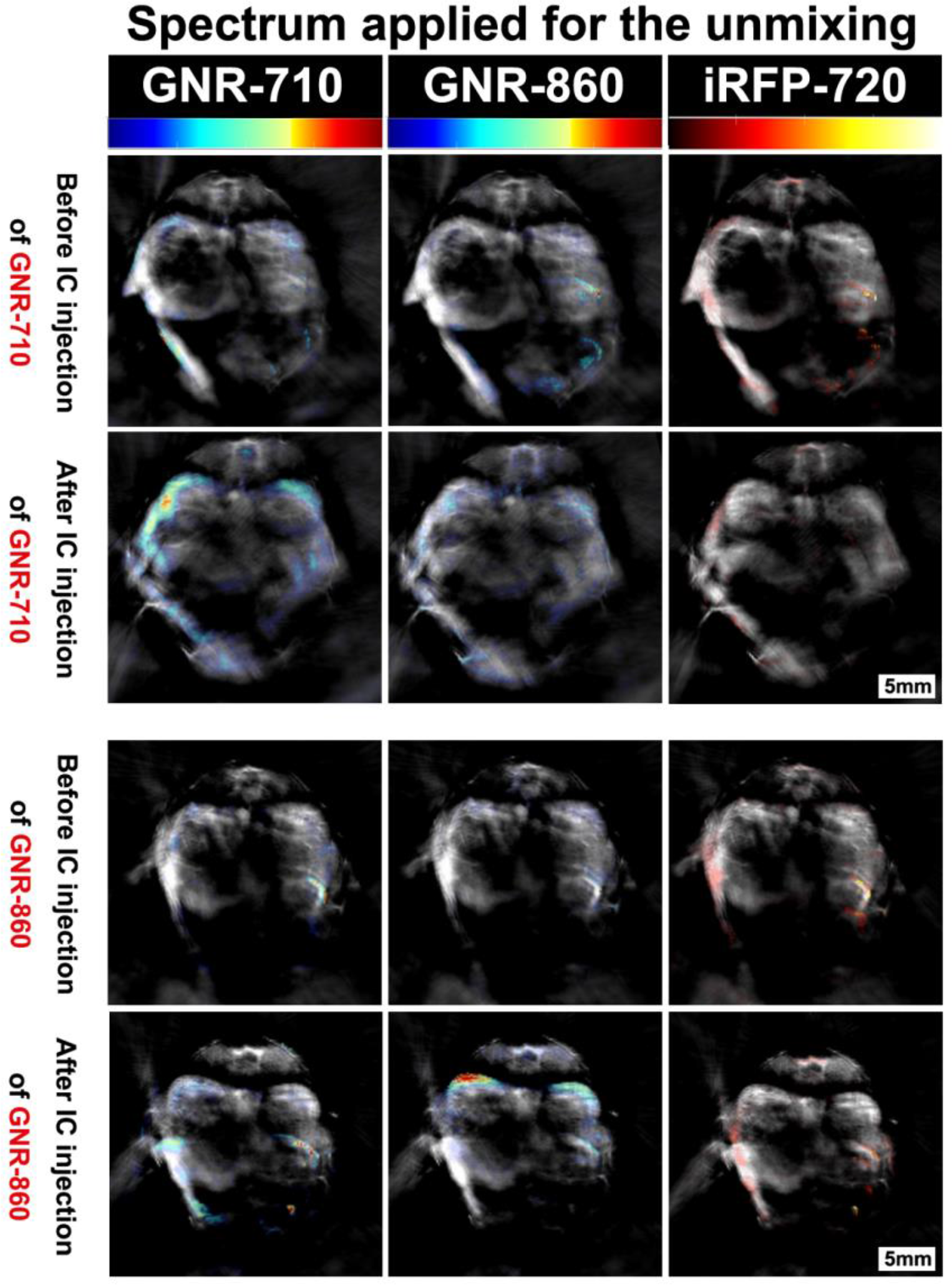
Specificity of the multispectral unmixing. Signal before and after treatment with GNR-710 labelled cells (upper panel) and GNR-860 labelled cells (bottom panel) in transverse sections at the level of the kidney region of the corresponding mouse. Signals were observed in the kidney only when the spectra of the respective component (GNR-710 in upper panel and GNR-860 in bottom panel) was applied. Note also that iRFP720 unmixing did not result in a signal at this stage. Colour scales are 0.1 to 14, 0.9 to 21, and 0.0 to 15 MSOT intensity (a.u.) for GNR-710, GNR-860, and iRFP720, respectively.

### Clearance and fate of the cells and GNRs after initial biodistribution

Within 24 hours after GNR-labelled cell infusion, only between 15-22% of the initial bioluminescence signal in whole mice persisted (Figure 5b), decreasing to less than 4% by day 5. Such substantial amounts of cell death following cell infusion were expected from previous reports ^40,41^. A similar degree of cell death was observed with cells not labelled with GNRs, indicating that cell loss *in vivo* was not caused by GNR labelling (Supplementary file 7). Interestingly, from this stage onwards the bioluminescence images no longer correlated with the photoacoustic imaging (Figure 5a-c), indicating that the vast majority of GNRs followed a different path than the remaining luciferase-expressing cells. Specifically, although the luminescence intensity had decreased substantially by 24 h, the distribution of the signal remained the same (Figure 5b). By contrast, the MSOT signal for GNR-860 increased dramatically in the liver over the same timescale, but was substantially reduced in regions such as the kidneys and head where it had been predominant immediately after cell injection (Figure 5a, c). Whilst luminescent signal is based on living cells, photoacoustic signal can be produced from GNRs independently, irrespective of whether they are within cells, or if the cells are viable. Thus, these data suggest that the liver plays an important role in the clearance of GNRs and potentially associated cell debris following the massive loss of cells during the first 24 h after injection, as previously reported ^42,43^. In addition, these findings are in line with reports describing nanoparticle clearance through the hepato-biliary tract^44–46^. We observed that the clearance could be continuously monitored in a longitudinal fashion since the photoacoustic signal in the liver dropped by 47% after 5 days and by 73% after 25 days (Figure 5d).

**Figure 5.**
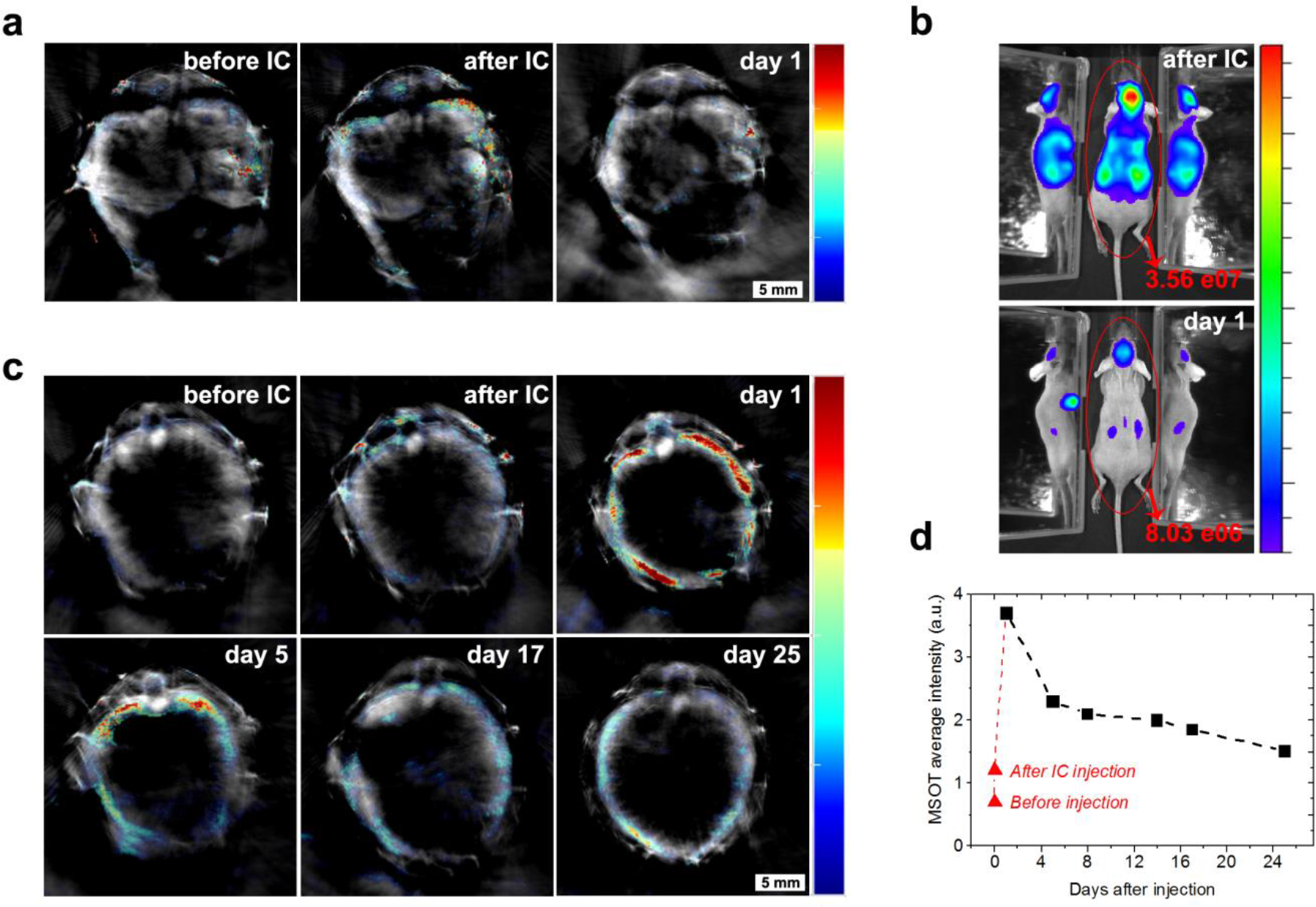
Cell death and clearance of GNR-860 from day 1 after injection. (a) MSOT signal in kidneys disappeared almost completely at day 1. (b) Total bioluminescence decreased by 78% at day 1 indicating a large amount of cell death. (c) In contrast to the kidneys, MSOT signal in liver increased substantially at day 1, which suggests that GNRs accumulated in the liver. (d) After day 1, MSOT signal in liver decreased progressively. Colour scales in (a) and (c) is 0.5 to 8 MSOT intensity (a.u.). Colour scale in (b) is 7.0 × 10^4^ to 5.8 ×10^5^ radiance units (ρ · s^−1^ · cm^−2^ · sr^−1^).

## Tumour monitoring

Following the substantial loss in cell viability over the first 24 h, signal intensity declined further and by day 5 cells were no longer detectable by BLI in the regions where they had first accumulated (i.e. brain, kidneys and spinal cord, Figure 3). However, we now observed BLI signals from cells that had settled in different locations (Supplementary File 8). The signal intensity increased over the following weeks as cells grew into tumours (Supplementary File 8). Since the mMSC cell line had been reported to form osteoid structures *in vivo* ^26^, and recent results from our group have revealed the capacity of the cells to form osteosarcomas ^47^, we anticipated the possibility of tumour development. In other studies, this process has been typically monitored by bioluminescence imaging ^48^, which allows the intensity of the signal to be correlated with the number of living cells. However, bioluminescence is limited by the poor spatial resolution at depths beyond 1 mm due to photon scattering. We therefore evaluated the potential of MSOT to determine the precise localisation of internal tumours as well as their size and shape. We used the constitutive expression of iRFP720 for the long-term monitoring of tumour growth by means of photoacoustic imaging. Presence of iRFP720 resulted in a change of the photoacoustic spectra in the regions where cells were forming tumours (Figure 6, Supplementary File 9). The fact that the iRFP720 absorption spectrum is very different from the main endogenous absorbers (a sharp band peaking at 690-700 nm, Supplementary File 9) facilitated its detection even with minimal changes of photoacoustic intensity with respect to the background.

Mice developed tumours in different locations whether the cells were labelled with GNRs or not (Figure 6 without GNRs and Figure 8 with GNRs). This process could be monitored longitudinally by bioluminescence and MSOT as shown in Figure 6, where a number of tumours formed in a mouse 30 days after administration of cells labelled with the iRFP720-Luciferase lentivirus. After scanning the animal in 1 mm steps, multispectral processing of the corresponding cross sections (Figure 6b) revealed the presence of tumours on the right shoulder (1), spinal cord (2), in the region dorsal of the kidney (2a and 2b), back (3), hip (4 and 5), and right leg (6). Tumours 2a and 2b are very illustrative of the outcomes that can be achieved with this approach. From the *in vivo* bioluminescence images (Figure 6a), it is difficult to determine the localisation of the tumours as either subcutaneous or in deep tissues, and with which organs they are associated. Analysis of photoacoustic images revealed that those tumours were growing in an unusual position close to and dorsal of the kidneys (Figure 6b). *Ex vivo* luminescence imaging confirmed the observations made with *in vivo* photoacoustic tomography (e.g. tumour masses 2a and 2b were found attached to the anterior-dorsal part of the kidneys) (Figure 6c). Expression of iRFP720 was also confirmed by *ex vivo* fluorescence imaging of these tumours (Supplementary File 10).

**Figure 6.**
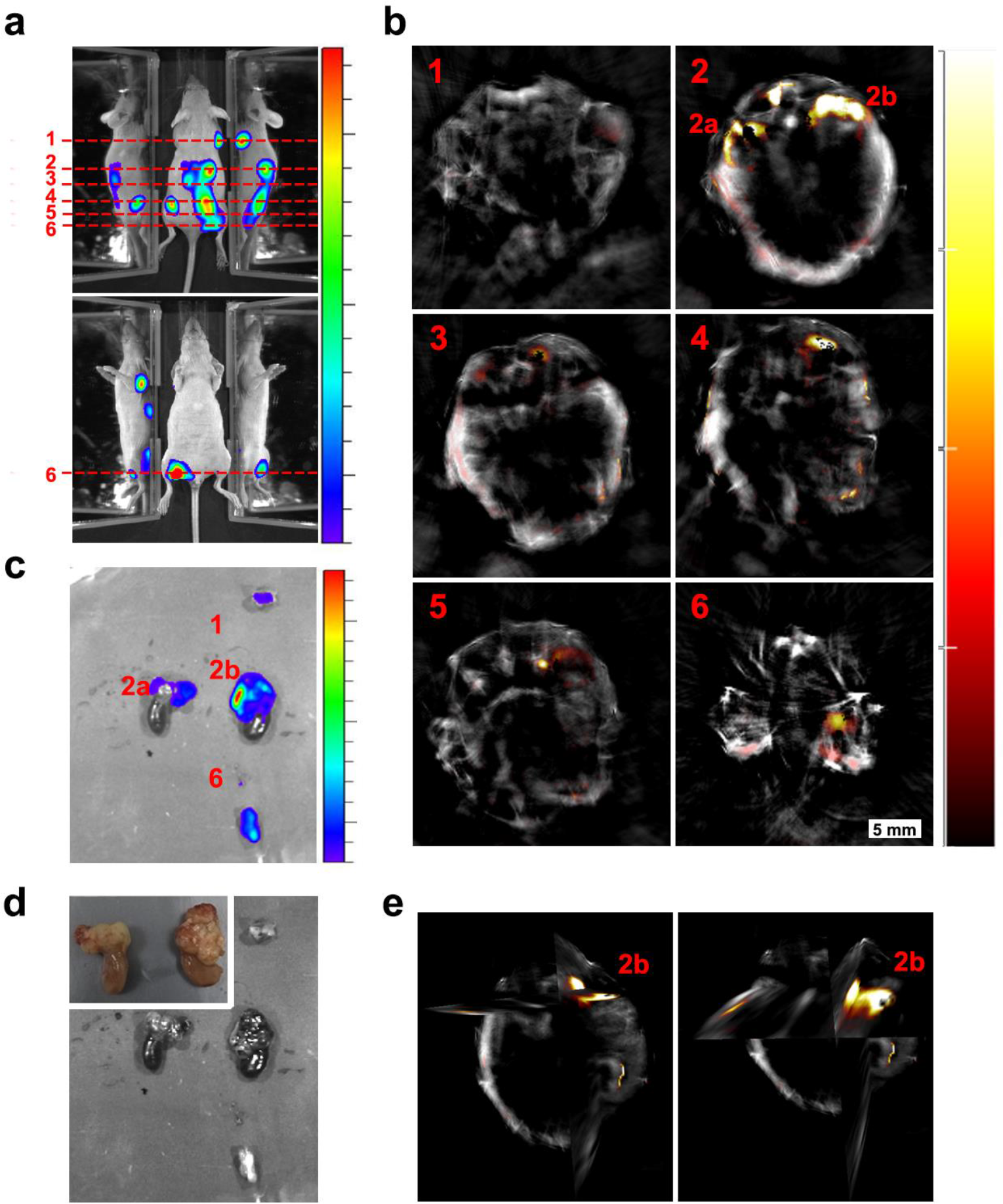
Localisation of tumours 30 days after injection of cells. (a) Bioluminescence images showing different tumours spread along the mouse body. Dashed lines indicate the approximate region shown in the MSOT scans. (b) Relevant MSOT transverse sections showing the precise localisation of different tumours after applying the multispectral processing for iRFP720. (c) *Ex vivo* luminescence imaging confirmed the localisation of relevant tumours imaged with MSOT. (d) *Ex vivo* picture of the tumours shown in (c) with a detail of kidneys in the inset. (e) Orthoslice images of tumour 2b in (b). Frontal view (left) and tilted view (right) provide a better spatial visualisation of the tumour′s precise localisation, extending anterior from the dorsal part of the right kidney. Colour scales are 1.4 × 10^8^ to 8.5 × 10^8^ radiance units (ρ · s^−1^ · cm^−2^ · sr^−1^) in (a) and 2.6 × 10^8^ to 5.3 × 10^9^ radiance units in (c). Colour scale in (b) is 0.4 to 15 MSOT intensity units (a.u.)

In addition to improving the anatomical localisation of internal tumours, photoacoustic tomography using iRFP720 expression also allowed precise longitudinal monitoring of tumour development from very early stages. The example in Figure 7 shows the growth of tumour 4 (Figure 6) from day 13 to day 30 after cell injection. The excellent spatial resolution enabled the localisation of this tumour in the vicinity of the pelvic ilium from day 13. The tumour mean diameter at this stage was 1.2 mm. By day 19, 23 and 30 it had expanded to 2.0, 2.2 and 3.3 mm, respectively. Furthermore, changes in the shape of the tumour were also monitored from day 23 onwards when it branched off into two ramifications, which grew attached to the main part of the tumour (Figure 7 and Supplementary File 11).

**Figure 7.**
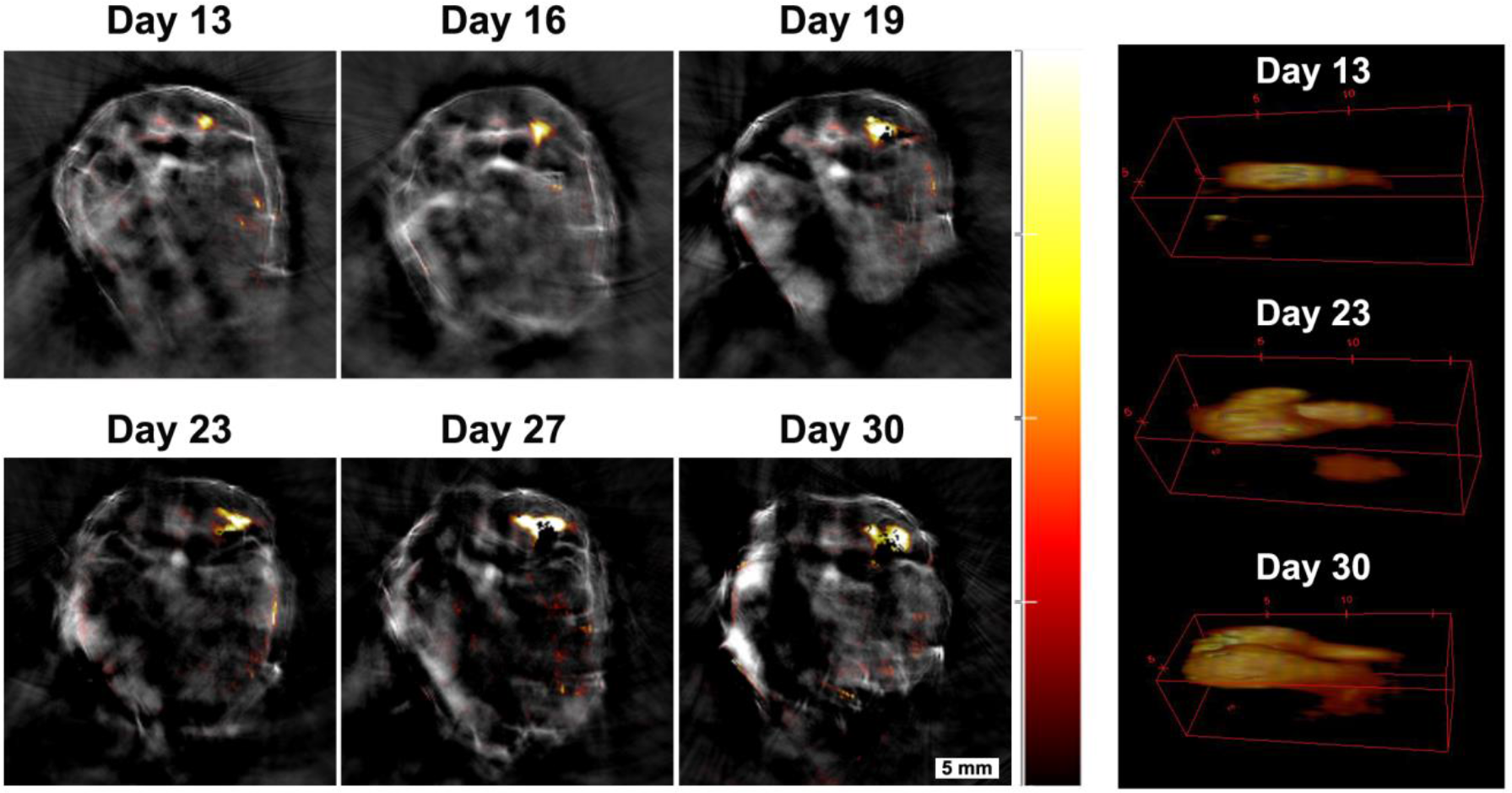
Monitoring tumour growth over time. Tumour 4 in Figure 6 was chosen to demonstrate the potential of the MSOT approach in monitoring tumour growth longitudinally over time. A small tumour was localised in the pelvic ilium region at day 13 and its growth was monitored from day 13 to day 30. 3D reconstructions of the tumour are shown on the right panel (360° animations are available as Supplementary File 11). Changes in size can be determined with this technique. Colour scale is 0.4 to 15 MSOT intensity units (a.u.)

In a second example, the presence of several tumours was determined 40 days after injection of GNR-860 and reporter gene labelled cells (Figure 8). Although GNRs were not used as contrast agents for tumour imaging, since the vast majority of them were cleared out following cell death, tumour growth was monitored with high spatial resolution via iRFP720 expression. The mouse shown in Figure 8 developed tumours in both shoulders, in the right dorsal kidney/adrenal gland region, towards the liver, and in several positions of the hip region. A late developing (days 33 – 40), but fast-growing tumour in the left shoulder was used to demonstrate the capabilities of iRFP720 MSOT imaging to obtain detailed spatial information in 3D reconstructions as shown in Figure 8c.

Overall, our analysis reveals that MSOT provides an excellent method to monitor tumour growth longitudinally from very early stages onwards (from sub mm size). Only some tumours growing in very peripheral positions of the animal could not be monitored due to technical or procedural limitations of the MSOT imaging system. For example, the nose could not be imaged as it was placed in the nose cone for air and anaesthesia supply during the imaging process. Also, since the animal was placed on its back, it was very difficult to avoid an air layer being trapped between the legs and the main body, which impaired sound propagation. Abundant use of ultrasound gel in this region minimised this effect and allowed data to be obtained as shown in Figure 6 (tumour 6), although the quality of the image was not as good as in other regions of the body. Any tumours in the lungs could not have been monitored with photoacoustic imaging due to a different sound propagation in air. However, the presence of tumours in the lungs could be ruled out from the bioluminescence images.

**Figure 8.**
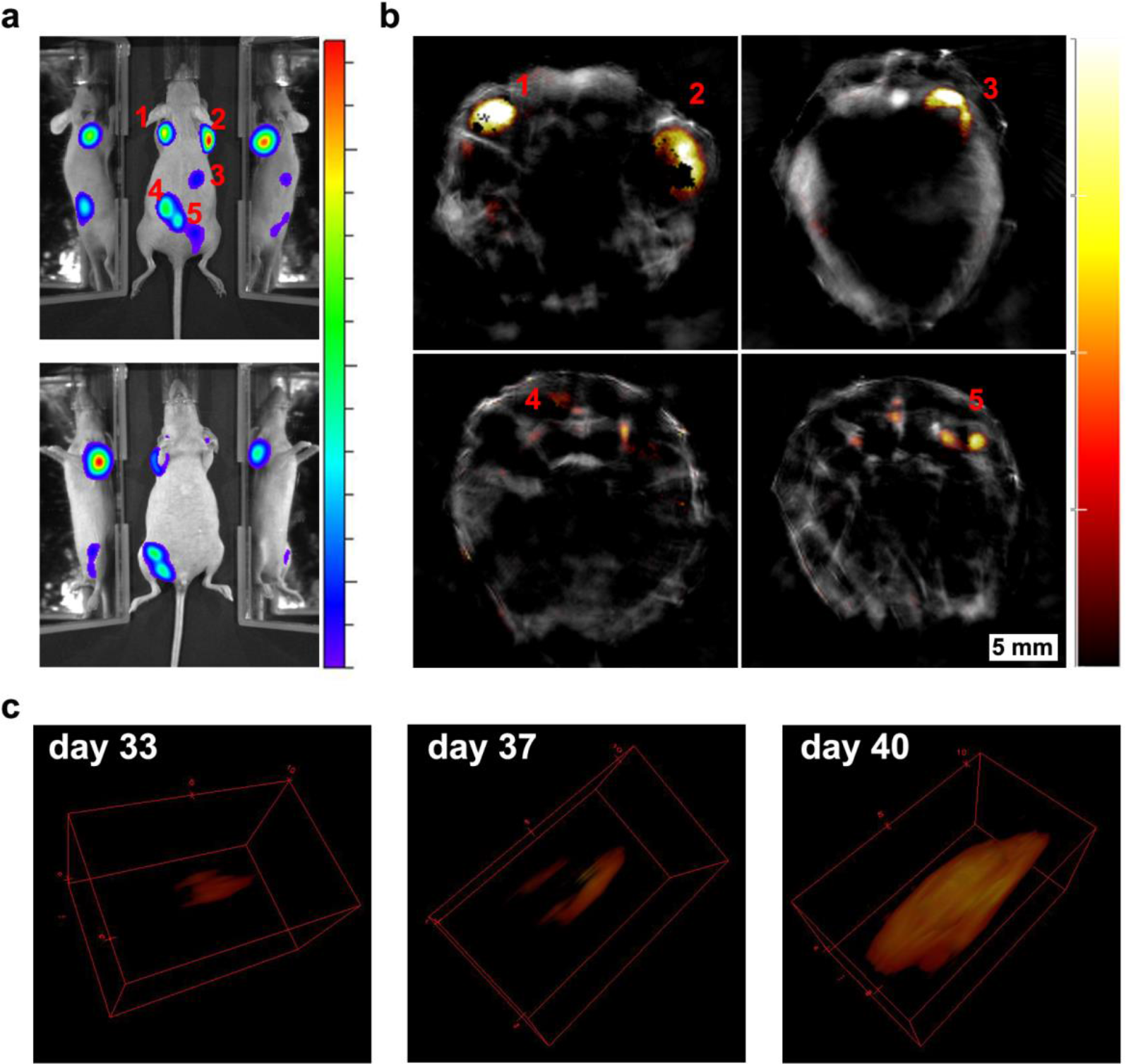
Longitudinal tumour monitoring in a mouse that was injected with GNR-860 / reporter gene labelled cells. Localisation of tumours 40 days after systemic injection of cells imaged by bioluminescence (a) and MSOT (b). (c) Growth of the tumour on the left shoulder was monitored from day 33 to day 40 and is shown in 3D reconstruction. Colour scale in (a) is 1.4 × 108 to 1.3 × 109 radiance units (ρ · s^−1^ · cm^−2^ · sr^−1^) and in (b) is 1.2 to 14 MSOT intensity units (a.u.). The black pixels visible within the tumour region in (b) are an experimental artefact (see Supplementary file 12 for additional discussion).

## Conclusions

We demonstrate here a multimodal imaging approach, which utilises a combination of GNRs and reporter genes to track cells immediately after systemic injection and longitudinally over time during the development of tumours. We show that multispectral optoacoustic tomography achieves high spatial resolution of the cell distribution without compromising sensitivity. Bioluminescence imaging, which is the standard optical-based imaging technology for *in vivo* cell tracking, was also performed for comparison and validation of our results.

Cells were labelled with silica-coated GNRs, which provided the excellent photoacoustic sensitivity required to monitor the broad distribution of cells after systemic intra-cardiac injection. Because the GNRs maintained their optical signature when taken up by the cells, spectral unmixing could be applied in MSOT, resulting in specific detection of labelled cells. Accumulation of cells in the brain, kidneys and liver was observed in agreement with bioluminescence imaging. Systemic administration of cells was followed by a high degree of cell death as indicated by a strong decline in bioluminescence and the separation of the photoacoustic GNR signal from the BLI signal of live cells. GNR signal became localised to the liver in line with its role in the hepato-biliary clearance of GNRs and associated cell debris.

To monitor the formation of tumours, the mMSCs had initially been modified with bi-cistronic reporter gene vectors to express constitutively both, firefly luciferase and iRFP720. Contrary to GNRs, the amount of iRFP720 increased with cell divisions, which made it an excellent candidate for photoacoustic monitoring of tumours. This approach allowed longitudinal tumour monitoring from very early stages (< 1 mm diameter). Precise anatomical localisation and morphology of tumours were evaluated using the superior deep tissue resolution of photoacoustic tomography compared to other optical-based imaging technologies.

Beyond the specific study of the biodistribution and tumour development of an exemplary mesenchymal cell line performed here, we emphasise the technological potential of these labelling tools for non-invasive *in vivo* imaging and cell tracking. In particular, research fields such as cell-based regenerative medicine therapies or cancer biology might find new applications using these labelling reagents and imaging approaches for cell monitoring and safety studies.

## Acknowledgements

We thank Joseph Zeguer and Danielle Vaughan for contributions to the lentivirus vector work, the Biomedical Services Unit, the Biomedical Electron Microscopy Unit and the Centre for Preclinical Imaging at The University of Liverpool for technical support, B. Kevin Park for helpful discussion on the direction of the project, MRC, BBSRC and EPSRC for sponsorship through the UK Regenerative Medicine Platform Safety Hub (MR/K026739/1), the EU FP7 and a Marie Curie Fellowship to JC (NANOSTEMCELLTRACKING).

## Supplementary material

**Supplementary file 1.**
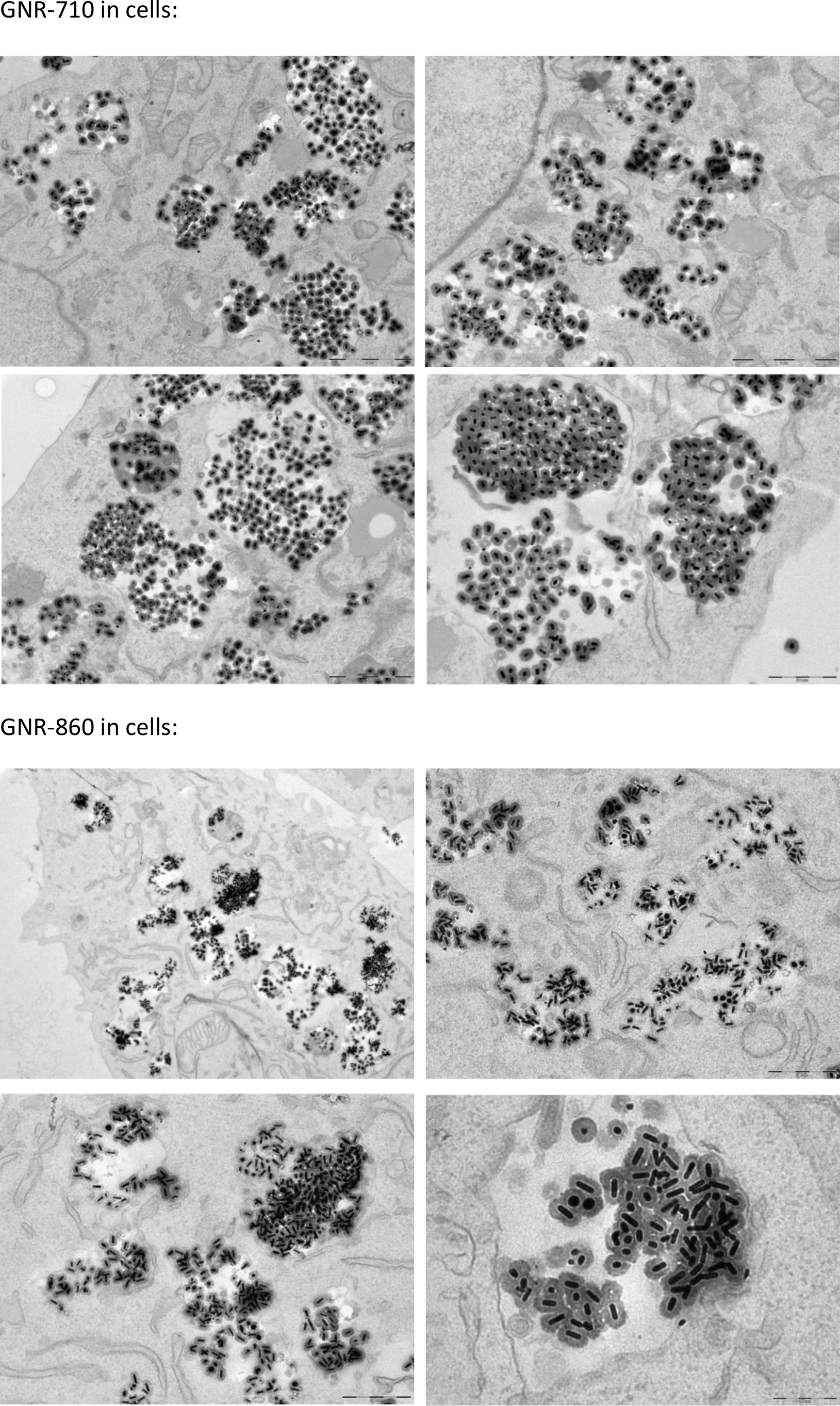
Additional images of GNR-710 and GNR-860 uptake by MSCs. Silica shell provided steric hindrance to minimise plasmon coupling effects.

**Supplementary file 2.**
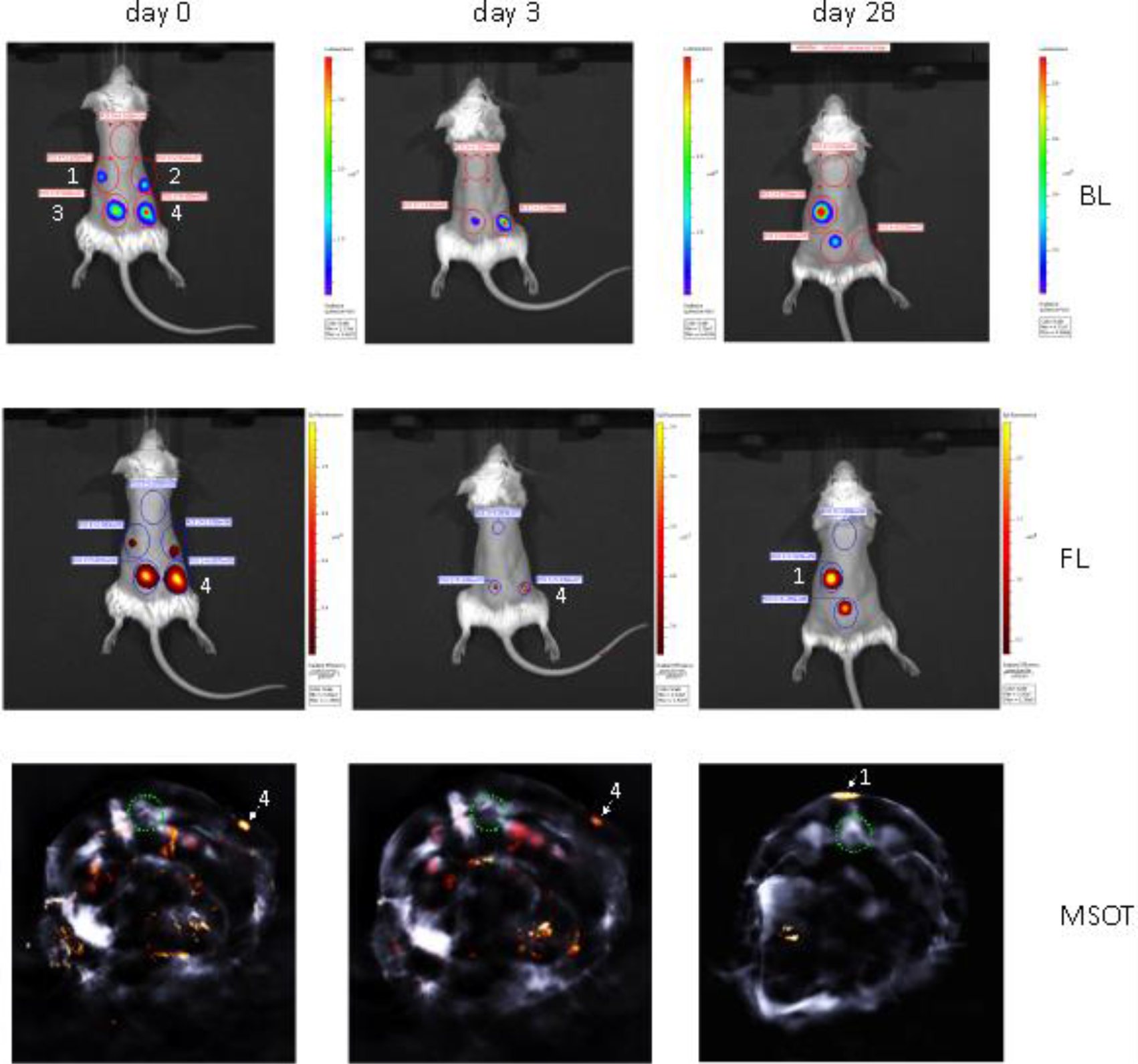
Longitudinal multi-modal imaging of iRFP720-E2A-Luc labelled cells after subcutaneous injection of mMSCs shows proportionality of bioluminescent, fluorescent and photoacoustic signals of the reporter genes. 1.25 × 10^5^ (1), 2.5 × 10^5^ (2), 5 × 10^5^ (3) and 1 × 10^6^ (4) cells were injected at the indicated regions and imaged immediately and over a period of one month after injection. A substantial loss of signal intensity over the first few days post-injection, which can be attributed to cell death, resulted in undetectability of the 1.25 × 10^5^ (1) and 2.5 × 10^5^ (2) cell injection clusters on day 3. However, survival of a small number of cells and subsequent tumour formation in the region of the lowest injected cell number (1) is evident after four weeks. Photoacoustic (MSOT) detection of iRFP720 is shown in tomographic transverse body sections at the site of highest cell concentration (1 × 10^6^, (4)) on the day of injection and on day 3, and for the larger tumour (1) on day 28. The photoacoustic signals are colour coded in arbitrary MSOT units and the green circle indicates the location of the spine for orientation. For iRFP720 fluorescence (FL) the 710/760 nm excitation/emission filter set was used. Exposure time was 1 second for both, FL and bioluminescence (BL). Multi-spectral data reconstruction for MSOT was performed employing a guided-ICA algorithm against the photoacoustic spectra of iRFP720-E2A-Luc cells in phantoms.

**Supplementary File 3. 360° animation of the 3D reconstruction shown in Figure 3 of the main text**

(supplementary file 3.avi)

**Supplementary file 4.**
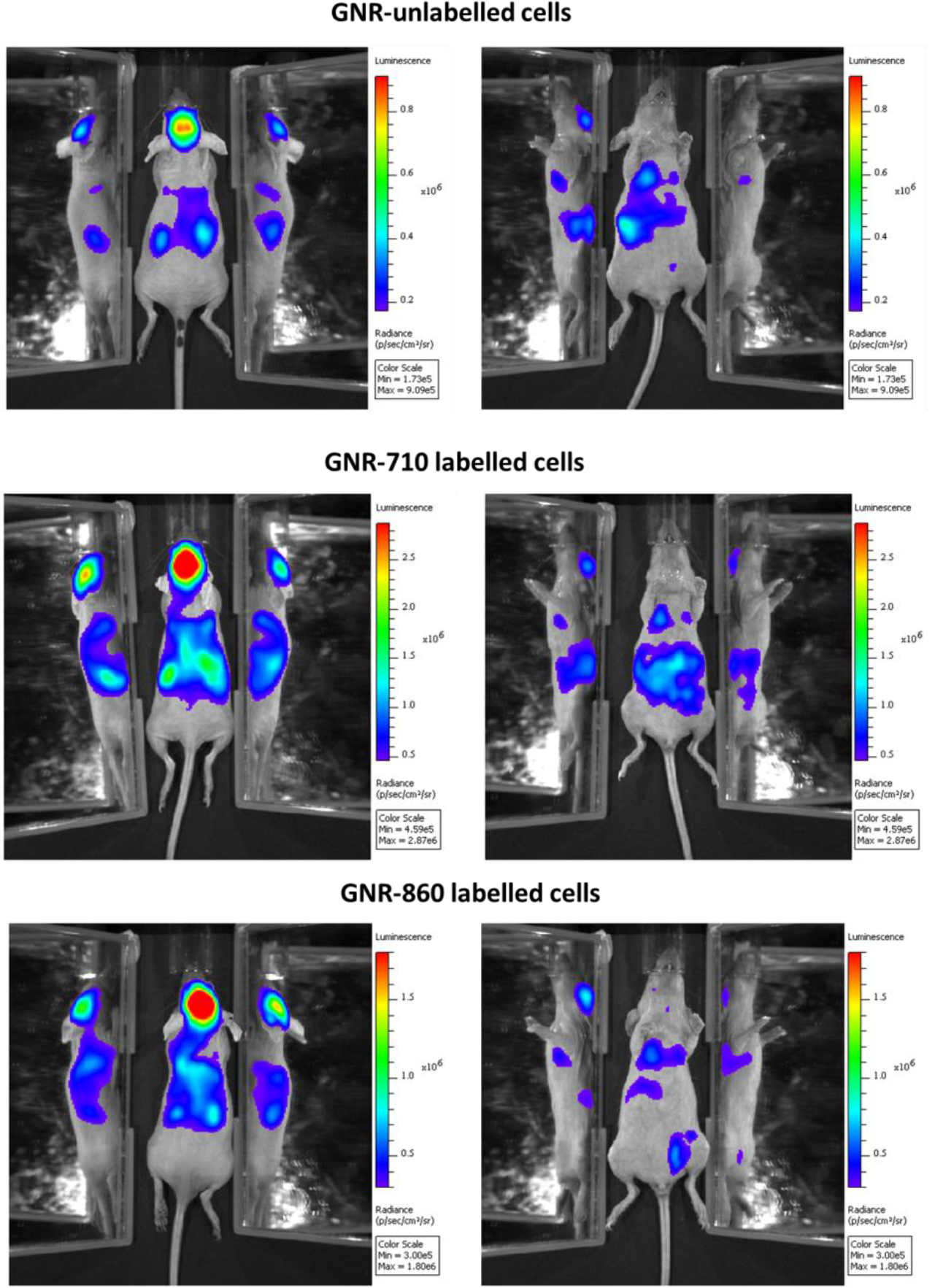
Biodistribution of cells not labelled with GNRs and cells labelled with GNR-710 and GNR-860, respectively, after intracardiac administration. A similar pattern was observed in all three cases, showing that GNRs did not affect overall cell distribution.

**Supplementary file 5.**
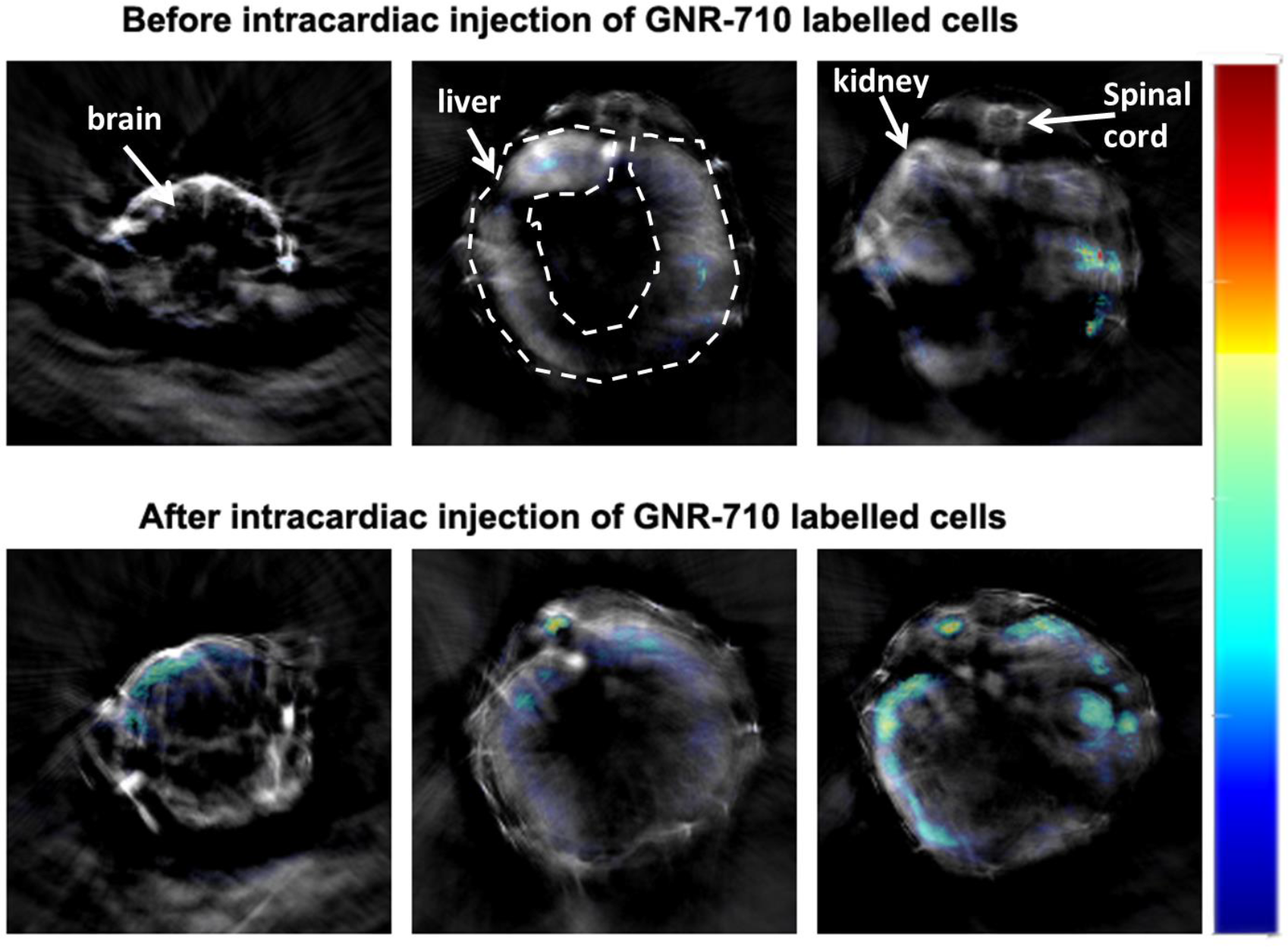
MSOT imaging of cell biodistribution of GNR-710 labelled cells. Transverse sections showing three regions of interest (brain, liver/spinal cord and kidneys) before and after the administration of cells. The results here are in agreement with the results for GNR-860 in the main text Figure 3. Colour scale is 0.8 to 25 MSOT intensity (a.u.). Note also that for GNR-710, we observed signal interferences in the intestines. This interference is likely due to a similar photoacoustic profile of food since this signal appeared also before the administration of cells, which helped to discriminate between interference and actual signal originating from GNRs.

**Supplementary file 6.**
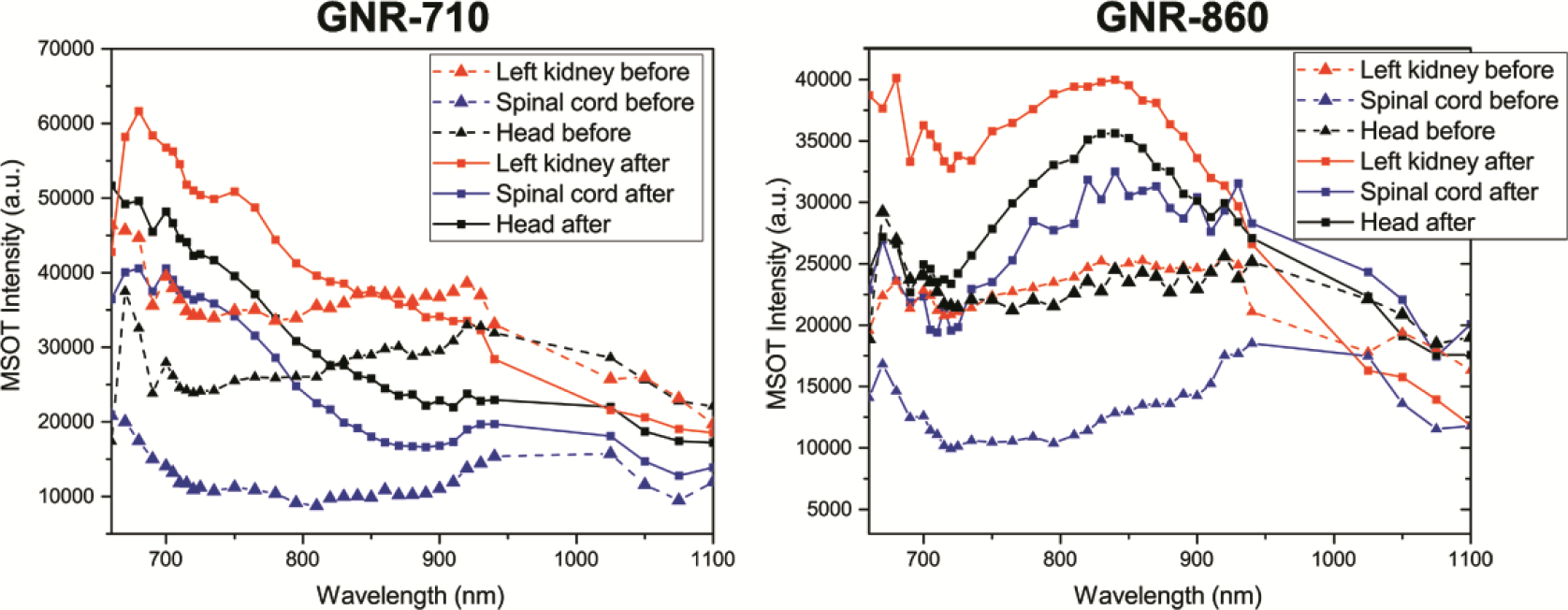
Photoacoustic spectra extracted from *in vivo* imaging. Photoacoustic spectra of three regions of interest (left kidney, spinal cord and head) before and after administration of GNR-710-labeled cells (left) and GNR-860-labeled cells (right). Note that the spectra corresponding to the scans before cell administration are similar in both cases, whilst they acquire the characteristic profile of their corresponding GNRs after cell administration. No wavelengths between 940 and 1025 were selected to avoid absorption by water, which has one absorption band in this region.

**Supplementary file 7.**
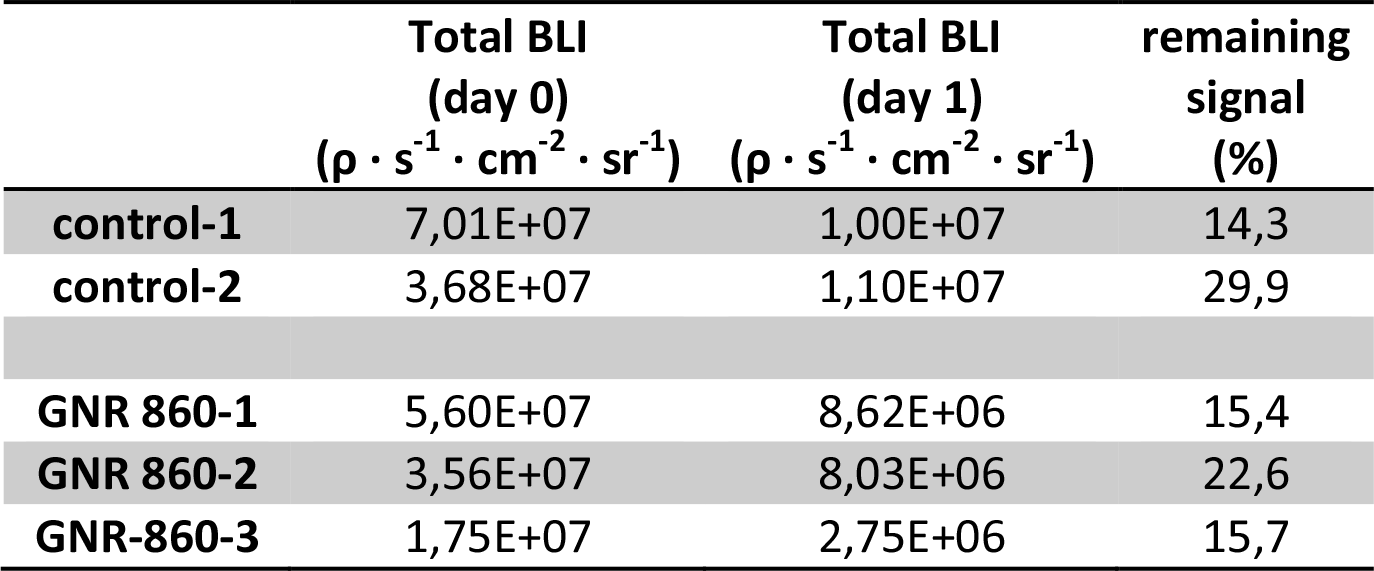
Loss in BLI signal intensity within 24 hours after injection. A similar percentage of decrease in total BLI signal intensity was observed in mice treated with GNR-unlabelled cells as in mice treated with GNR-860-labelled cells, suggesting that GNRs did not affect cell death.

**Supplementary file 8.**
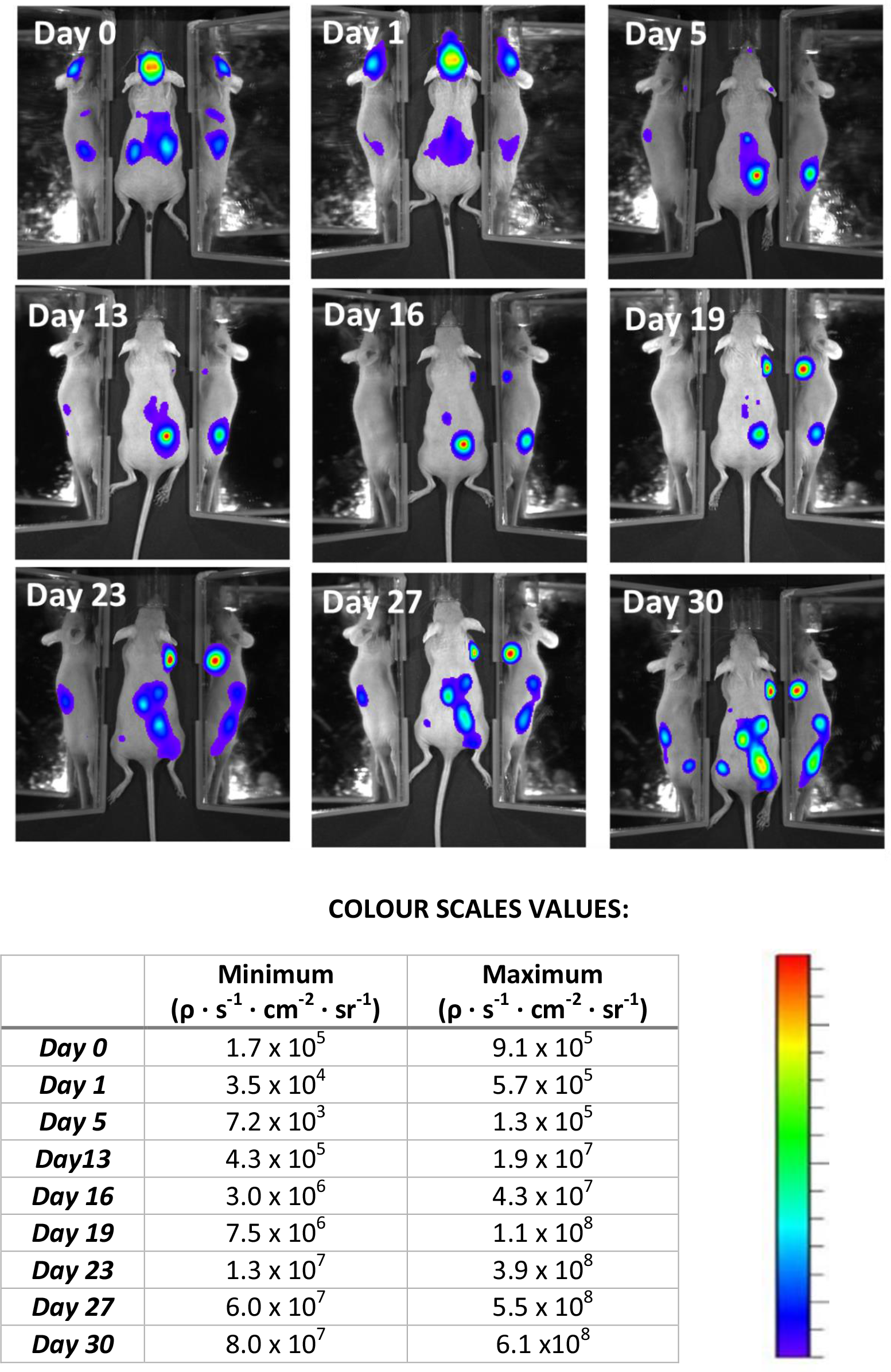
Bioluminescence monitoring of tumour growth of the mouse shown in Figures 6 and 7 of the main text. Several tumours developed and became detectable from five days after cell injection. Colour scales have been adjusted to show tumour localisation. They cannot be standardized here due to the great increase of intensity after tumour growth.

**Supplementary file 9.**
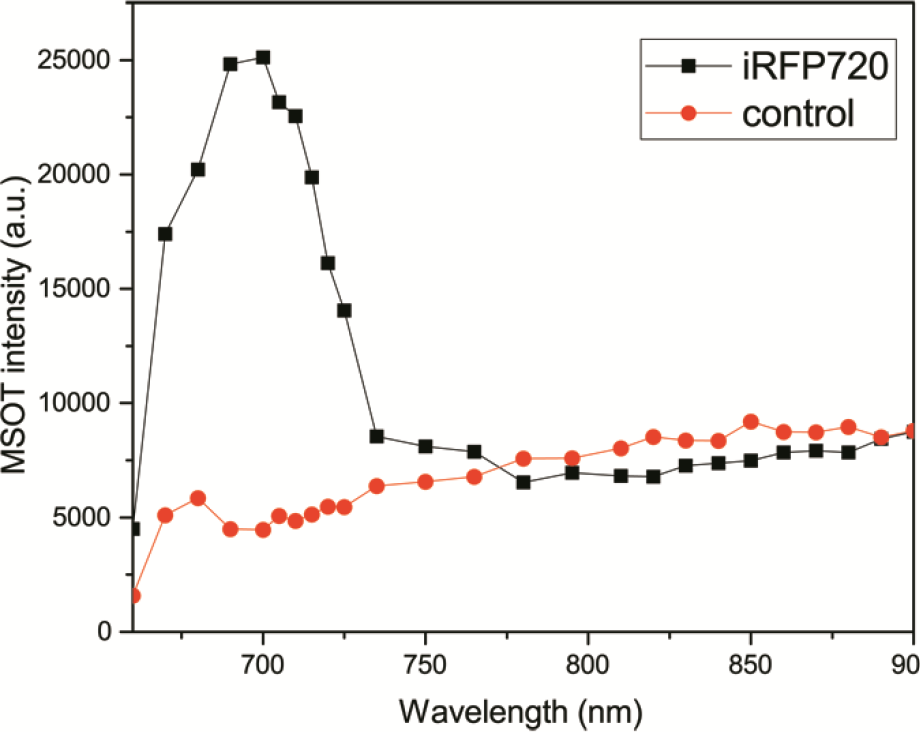
*In vivo* photoacoustic spectra of tumour 2b in Figure 6 of the main text. The characteristic spectrum of iRFP720 is obtained after analysing the photoacoustic intensities at the working wavelengths. A random region below the tumour was chosen as a control region for comparison.

**Supplementary file 10.**
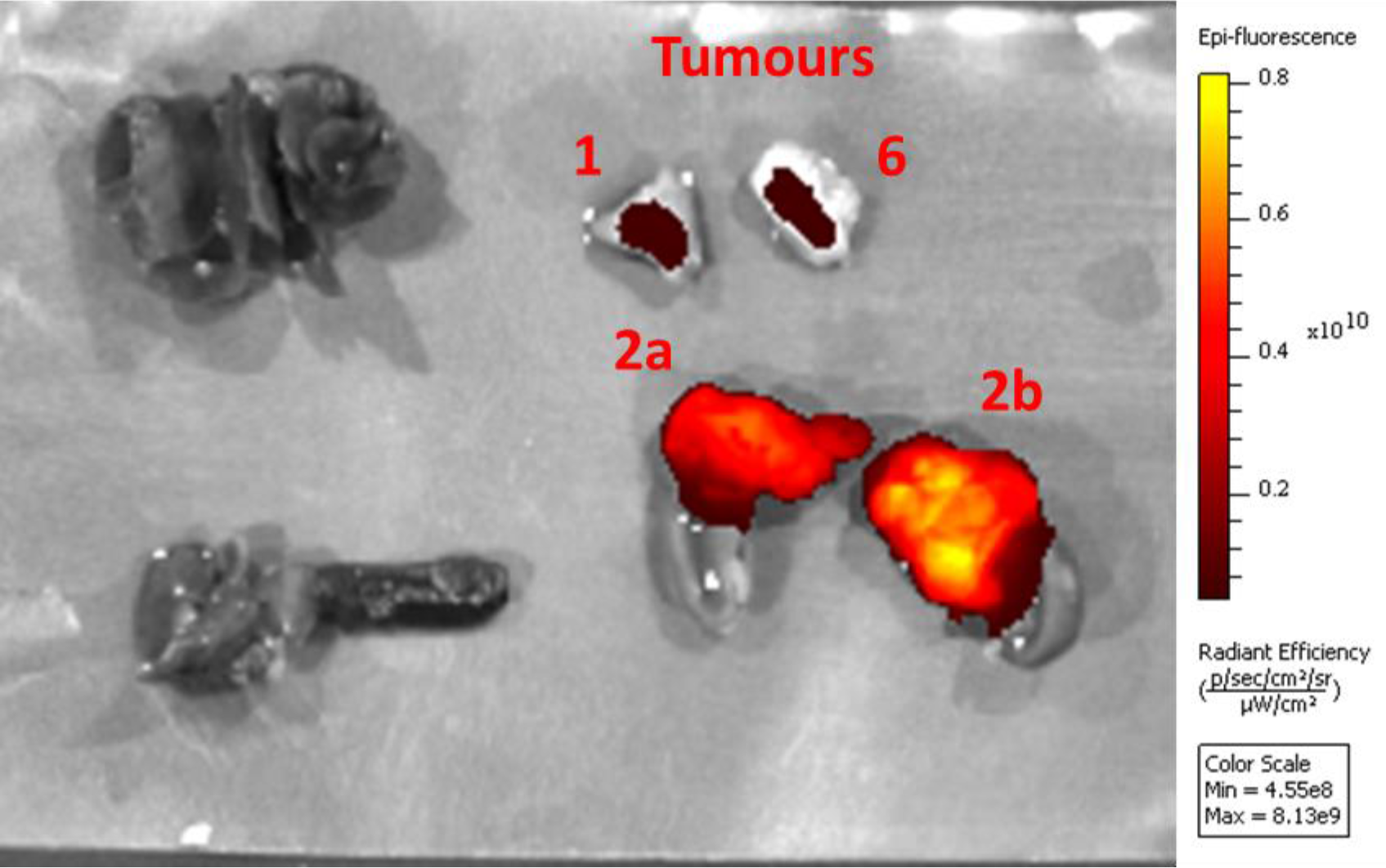
*Ex vivo* epi-fluorescence image (Ex=675 nm, Em=720 nm) of tumours shown in Figure 6, after fixation with 4% PFA for 24 hours. No signal was observed in organs such as liver (top left), stomach and attached spleen (bottom left), and heart and lungs, indicating the specificity of the signal.

**Supplementary File 11. 360° animation of the 3D reconstructions shown in Figure 7 of the main text.**

(supplementary file 11.pptx)

**Supplementary file 12.**
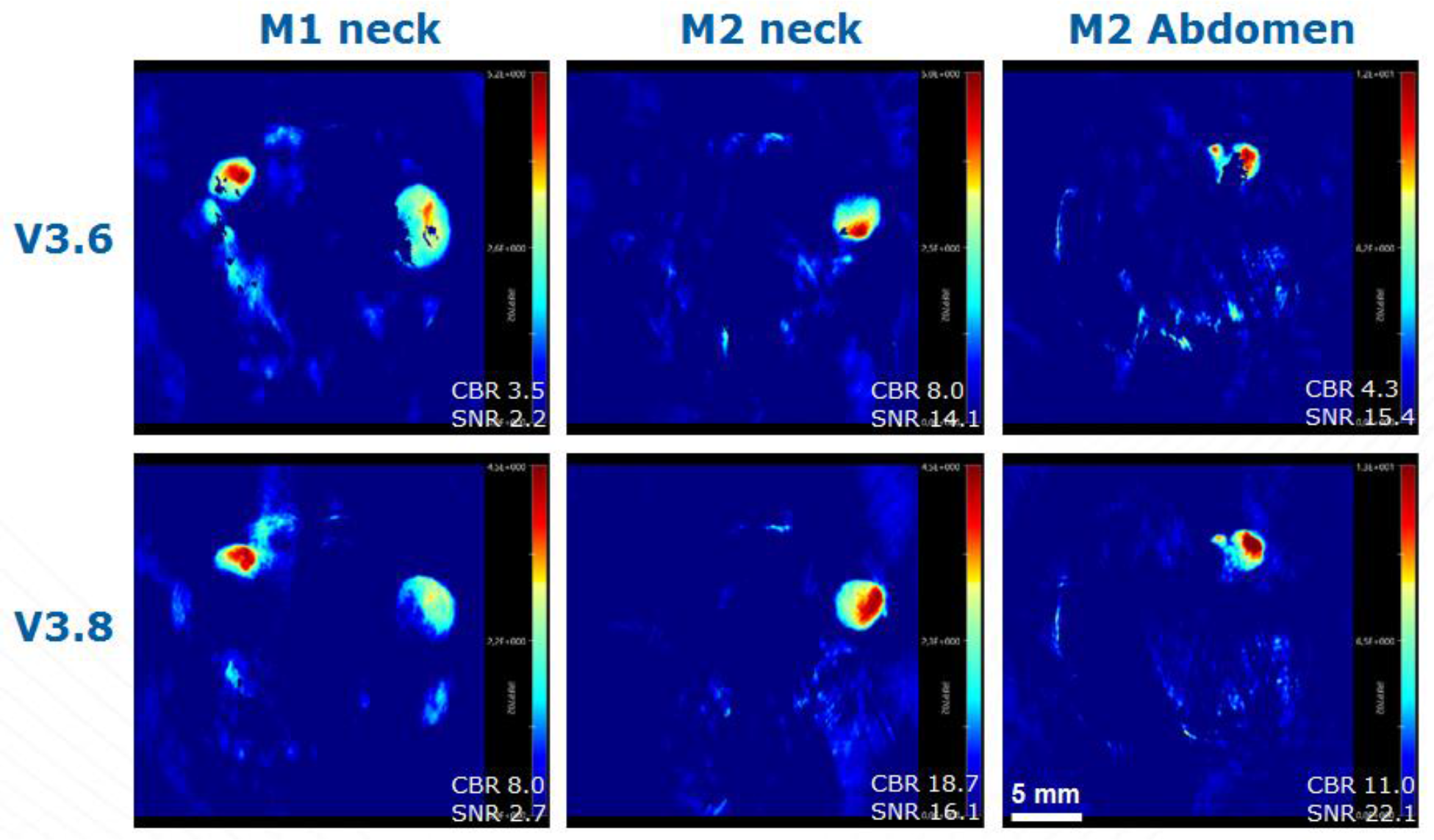
The negative areas in the reconstructed MSOT images (appearing as black pixels in some of our figures) have a variety of origins. For all of the figures of this article, v3.6 of the MSOT software was used. v3.8 has now been released. It includes a better model for the receive characteristics of the transducer that reduces the number of negative pixels as can be seen above. Thank you to Stefan Morscher and Thomas Sardella of iTheraMedical for helpful discussion and providing the figure above. M1 is mouse GNR-860 (group III in table 1 of the main ms) and M2 is a control mouse (group I in table 1 of the main ms).

